# Reptile-like physiology in Early Jurassic stem-mammals

**DOI:** 10.1101/785360

**Authors:** Elis Newham, Pamela G. Gill, Philippa Brewer, Michael J. Benton, Vincent Fernandez, Neil J. Gostling, David Haberthür, Jukka Jernvall, Tuomas Kankanpää, Aki Kallonen, Charles Navarro, Alexandra Pacureanu, Berit Zeller-Plumhoff, Kelly Richards, Kate Robson-Brown, Philipp Schneider, Heikki Suhonen, Paul Tafforeau, Katherine Williams, Ian J. Corfe

## Abstract

There is uncertainty regarding the timing and fossil species in which mammalian endothermy arose, with few studies of stem-mammals on key aspects of endothermy such as basal or maximum metabolic rates, or placing them in the context of living vertebrate metabolic ranges. Synchrotron X-ray imaging of incremental tooth cementum shows two Early Jurassic stem-mammals, *Morganucodon* and *Kuehneotherium*, had lifespans (a basal metabolic rate proxy) considerably longer than comparably sized living mammals, but similar to reptiles. *Morganucodon* also had femoral blood flow rates (a maximum metabolic rate proxy) intermediate between living mammals and reptiles. This shows maximum metabolic rates increased evolutionarily before basal rates, and that contrary to previous suggestions of a Triassic origin, Early Jurassic stem-mammals lacked the endothermic metabolism of living mammals.

**One Sentence Summary:** Surprisingly long lifespans and low femoral blood flow suggest reptile-like physiology in key Early Jurassic stem-mammals.

## Main Text

Recent discoveries and analyses have revolutionized our knowledge of Mesozoic mammals, revealing novel aspects of their ecology *(1, 2)* development *(3, 4)* systematics *(2, 4)* and macroevolution *(5, 6)*. However, details of physiology are more difficult to determine from fossils, and our knowledge of physiological evolution remains comparatively poor. Living mammals are endotherms, possessing the ability to control and maintain metabolically produced heat, and have a substantially higher capacity for sustained aerobic activity than ectothermic animals *(7-9)*. The origin of endothermy is an important event in mammalian evolution, often noted as key to their success *(7-9)*. Several competing hypotheses seek to explain the selective pressures and adaptive pathways of endothermic evolution: (a) selection for higher maximum metabolic rates (MMR) enhanced sustained aerobic activity *(7, 10, 11)*; (b) selection for higher basal metabolic rates (BMR) enhanced thermoregulatory control *(12, 13)*; or (c) MMR and BMR evolved in lockstep with each other *(8, 9)*.

Direct evidence from living mammals to support these hypotheses is equivocal *(7)*. Recent analyses find no long-term evolutionary trend in BMR *(14)* contradicting earlier suggestions of increasing BMR throughout the Cenozoic *(13)*, and so implying that the Middle Jurassic (∼170 Ma) most recent common ancestor (MRCA) of living mammals *(14)* possessed a BMR within the range of present-day mammals. Several indirect indicators of metabolic physiology in fossil synapsids have been suggested but provide contradictory evidence for the timing of origin of endothermy and its evolutionary tempo. These include: the presence of fibrolamellar long-bone, first seen in the Early Permian (∼300 Ma) synapsid *Ophiacodon (15)*; the presence of an infraorbital canal and lack of parietal foramen, used to infer facial whiskers, fur, lactation and endothermy in Early Triassic (∼245 Ma) cynodonts *(16)*; inferred maxillary nasal turbinates in the Late Permian (∼255 Ma) therapsid *Glanosuchus*, used to suggest that mammalian levels of endothermy evolved by the Late Triassic (∼210 Ma) *(17)*; a trend toward increased relative brain size initiated in Late Triassic non-mammaliaform cynodonts *(18)* and the mammaliaform stem-mammal *Morganucodon (19, 20)*; and acquisition of a parasagittal gait in the Early Cretaceous (∼125 Ma) therian mammal *Eomaia (21)*. Several recent studies provide more quantitative links to physiological parameters. Oxygen isotopes were used to infer elevated thermometabolism in Middle-Late Permian (∼270-255 Ma) eucynodonts *(22)*; red blood cell size diminution in Late Permian (∼255 Ma) eutheriodontid therapsids was qualitatively linked via two proxies to increased MMR *(23)*; and osteocyte lacuna shape correlations suggested ‘mammalian’ resting metabolic rates in Permo-Triassic (∼250 Ma) dicynodonts *(24)*.

However, the inconsistency of these characters, in time and with respect to phylogeny *(25, 26)*, along with re-assessments of function in relation to endothermy *(8, 27)*, limit their use as conclusive indicators of modern mammalian levels of endothermy in fossil taxa. Such temporal and phylogenetic heterogeneity suggests the evolution of mammalian endothermy followed a complex, mosaic pattern with different physiological aspects evolving independently, and at separate rates, towards current mammalian levels. Additionally, few of these physiological proxies are directly related to measurable aspects of metabolic rate.

To address these issues, we used two proxies to improve understanding of physiology at one of the most important nodes along this transition by estimating BMR and calculating a known proxy for MMR for two of the earliest mammaliaforms, *Morganucodon* and *Kuehneotherium (1, 28)*.

### Lifespan: a proxy for mammaliaform physiology

We used maximum lifespan estimates for fossil mammaliaform taxa as a proxy for BMR *(29)*. In extant tetrapods, negative correlations exist between lifespan and BMR *(29)* (and lifespan and growth rate *(30)*, another indicator of metabolism - Supplementary Materials). In general, the longer a mammal’s lifespan, the lower its size-adjusted BMR, so an accurate assessment of mammaliaform lifespan can be used to estimate metabolic potential. To do so we used cementochronology, which counts annual growth increments in tooth root cementum, and has been widely used to record lifespans in extant mammals *(31, 32)*. Cementum is a mineralized dental tissue surrounding the tooth root (Fig 1A, B), attaching it to the periodontal ligament and anchoring the tooth within the alveolus *(31)*. Cementum growth is continuous throughout life in extant mammals and seasonally appositional in nature, forming a series of increments of differing thickness and opacity when viewed in histological thin-sections under light microscopy. The correlation between increment count and chronological age is well documented, with one thick and one thin increment deposited every year *(31)*, where the thin, hyper-mineralized opaque increments record growth rate reduction in less favourable seasons *(33)*.

**Fig. 1.**
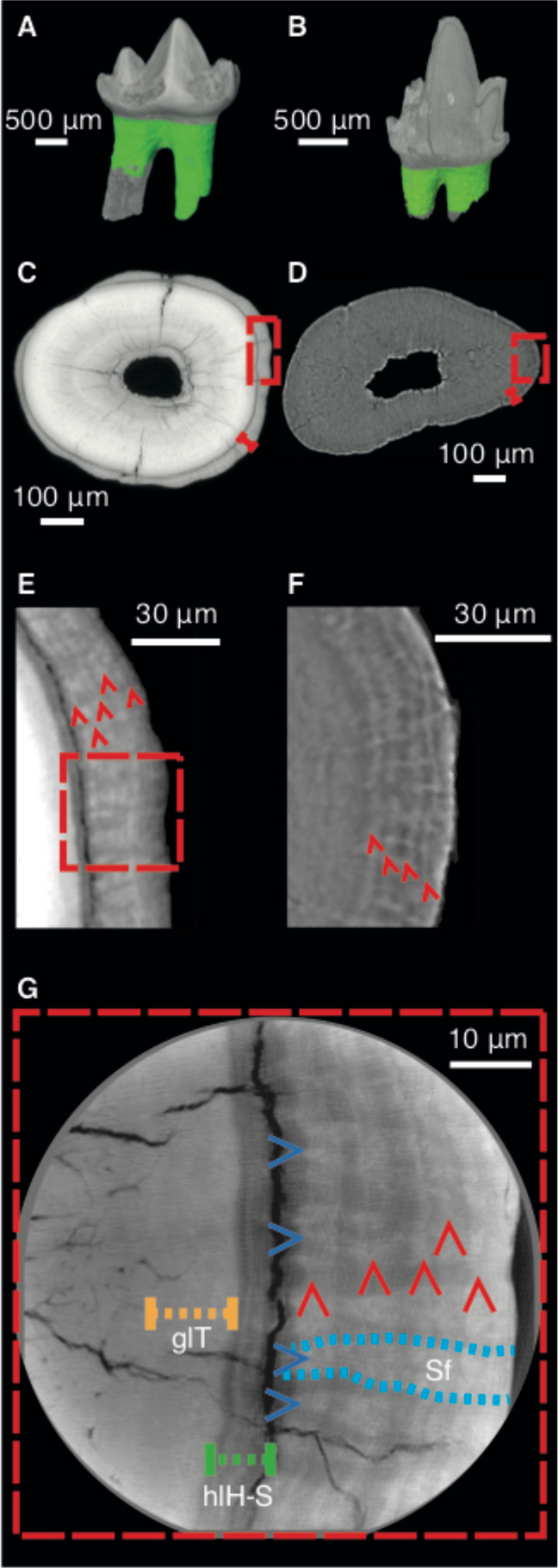
Cementum of *Morganucodon* and *Kuehneotherium*. **A**, **B.** 3D reconstructions of **A**, *Morganucodon* right lower molar tooth NHMUK PV M 104134 (voxel size 2 μm, μCT) and **B**, *Kuehneotherium* right lower molar tooth NHMUK PV M 21095 (voxel size 1.2 μm, PPC-SRµCT). Green = cementum. **C**, **D.** Transverse PPC-SRµCT virtual thin-sections (0.33 μm voxel size) of roots of **C**, NHMUK PV M 104134 and **D**, NHMUK PV M 27436 (*Kuehneotherium*). Red bracketed line highlights cementum surrounding dentine. **E**, **F.** Close-ups of boxes in **C**, **D**, with five and four circumferential light/dark increment pairs highlighted by red arrows, respectively. **G**. Synchrotron nanotomographic virtual thin-section of *Morganucodon* (30 nm voxel size) - close-up of region near box in **E.** Vertical red arrows = cementum increments; horizontal blue arrows, dashed blue lines and Sf = radial bands of Sharpey’s fibres; yellow dashed line and glT = granular layer of Tomes; green dashed line and hlH-S = hyaline layer of Hopewell-Smith.

Despite this potential, cementochronology has not previously been attempted for fossil mammals older than the Pleistocene (2.6 Ma) *(34)*, because histological thin-sections destroy fossils and provide only a restricted field-of-view. We overcame these problems by using propagation phase-contrast X-ray synchrotron radiation microtomography (PPC-SRµCT) to non-destructively image fossilized cementum increments *(35, 36)*. The sub-micrometre resolution, fast-throughput and three-dimensional (3D) nature of PPC-SRµCT allows large sample sizes and volumetric datasets of increments along their entire transverse and longitudinal trajectories. Cementum increments are known for lensing and coalescence, creating errors in counts based on single, or limited numbers of, two-dimensional thin sections created for each tooth *(31)* (Fig S1). PPC-SRµCT imaging and 3D segmentation of individual cementum increments across extensive vertical distances allows principal annual increments to be distinguished from accessory increments created by lensing and coalescence (Materials and Methods).

*Morganucodon* and *Kuehneotherium* are coexisting Early Jurassic (∼200 Ma) shrew-sized insectivores1, known from thousands of teeth and bones (Materials and Methods), providing a rare opportunity to analyse the population-sized samples needed for confident maximum lifespan estimation. Importantly, these are the earliest diphyodont taxa (Fig S2), with a single replacement of non-molar teeth and no molar tooth replacement *(28)*, and so estimates of lifespan are accurate to the time of the measured tooth root formation. We applied PPC-SRµCT to isolated teeth, and mandibles with multiple teeth or roots *in-situ*, totalling 87 *Morganucodon* specimens (52 isolated teeth, 35 dentaries), and 119 *Kuehneotherium* specimens (116 isolated teeth, 3 dentaries). From these, 34 *Morganucodon* and 27 *Kuehneotherium* specimens were sufficiently preserved for three observers to independently estimate lifespan from cementum increments and compare for accuracy and precision validation (Table S1). The remainder showed physical and/or diagenetic damage preventing increment counting (Fig S3).

The cementum of *Morganucodon* and *Kuehneotherium* (Fig 1A, B) is distinguished from dentine in our PPC-SRµCT data by a distinct boundary layer, which lies external to the dentine’s granular layer of Tomes and is interpreted as the hyaline layer of Hopewell-Smith (Fig 1C-G). Synchrotron nanotomographic imaging (30 nm isotropic voxel size) of several exceptionally preserved specimens highlights individual Sharpey’s fibre bundles (linking cementum to the periodontal ligament in extant mammals), extending radially through the cementum (Fig 1G). In transverse section, cementum is ∼10-70 μm radially thick and displays contrasting light and dark circumferential increments representing different material densities (Figs 1E-G, 2A-D). Higher density increments (represented by greater greyscale values) average 2-3 μm radial thickness, and lower density increments 1-3 μm (Figs 1C-G, 2A-D). Individual increments can be followed continuously longitudinally and transversely through the entire tooth root (Fig 2E, F).

**Fig. 2.**
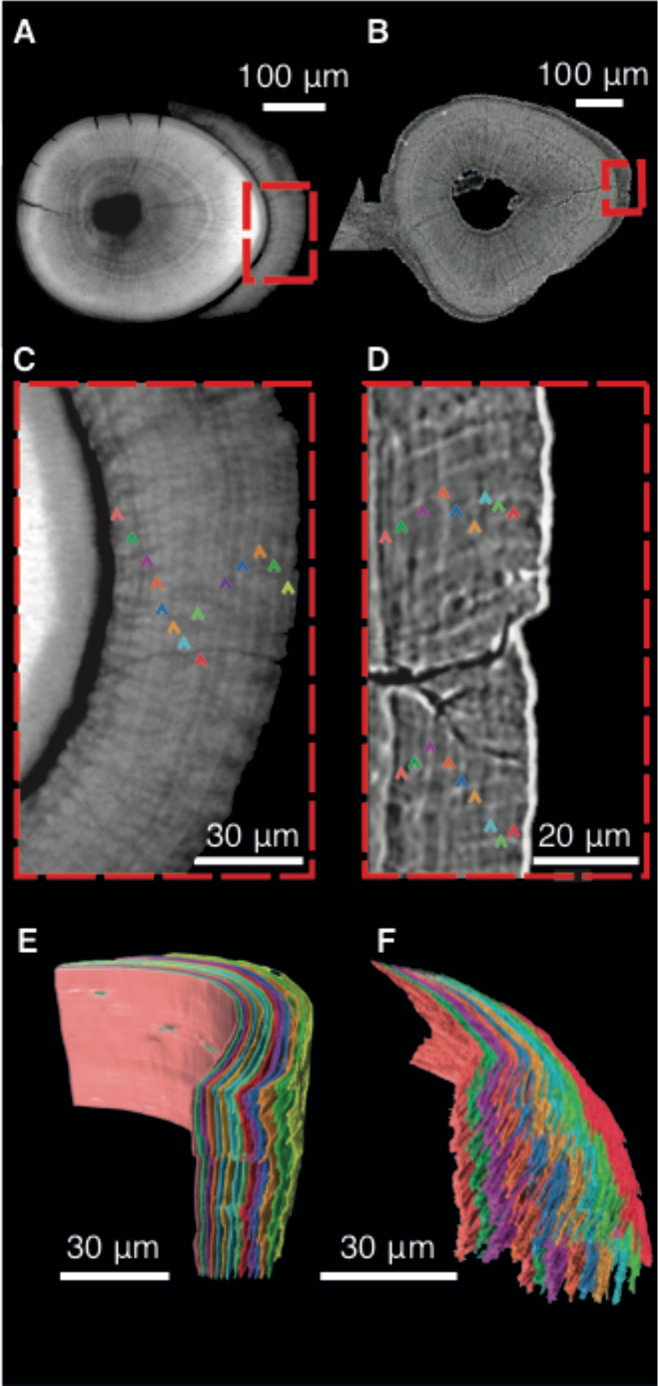
Three-dimensional segmentation of *Morganucodon* and *Kuehneotherium* specimens with the highest cementum increment counts. **A**, **B.** Transverse virtual thin-sections of PPC-SRµCT reconstructions (0.33 μm^3^ voxel size). **A**, *Morganucodon* specimen NHMUK PV M 104127 showing 55 μm thick cementum layer around root dentine. **B**, *Kuehneotherium* specimen UMZC Sy 141 showing 32 μm thick cementum. **C**, **D.** Detail of cementum of **C**, *Morganucodon* and **D**, *Kuehneotherium*. Cementum increments highlighted by 14 and nine multi-colouredarrows, respectively. **E**, **F.** 3D segmentations of cementum increments of **E**, *Morganucodon*, and **F**, *Kuehneotherium*. The colour of each increment corresponds to arrow colour in **C**, **D**.

We tested the accuracy of cementum increment counts for predicting mammaliaform lifespan by additionally PPC-SRµCT imaging eight dentulous *Morganucodon* specimens with a range of teeth *in-situ*. We counted cementum increments for several teeth along the tooth rows, and lines of arrested growth (LAGs) in the dentary bone periosteal region in two of these (Fig 3). Comparisons between counts of cementum increments are identical across all four premolars (p1-p4) and the anterior-most molars m1 and m2 in all specimens where they occur together (Table S2). Counts are also identical between dentary LAGs and cementum increments in the teeth present, p3-m2, of both specimens where LAGs were found (Fig 3). In three specimens with m1-3, m3 has one less increment than m1/m2. In one specimen i4 and the canine have one less increment than p1–3. This agreement between p1-m2 teeth and dentary increment counts indicates growth in both teeth and jaws was following the same, circum-annual rhythm previously reported for multiple extant mammal species *(31)*. We consider this strong support for the accuracy of lifespan estimates based on these increment counts. Additionally, the differences in increment counts along toothrows show eruption of the adult dentition in *Morganucodon* occurred over more than one year, which is considerably slower than in extant small mammals *(31)* (Supplementary text).

**Fig. 3.**
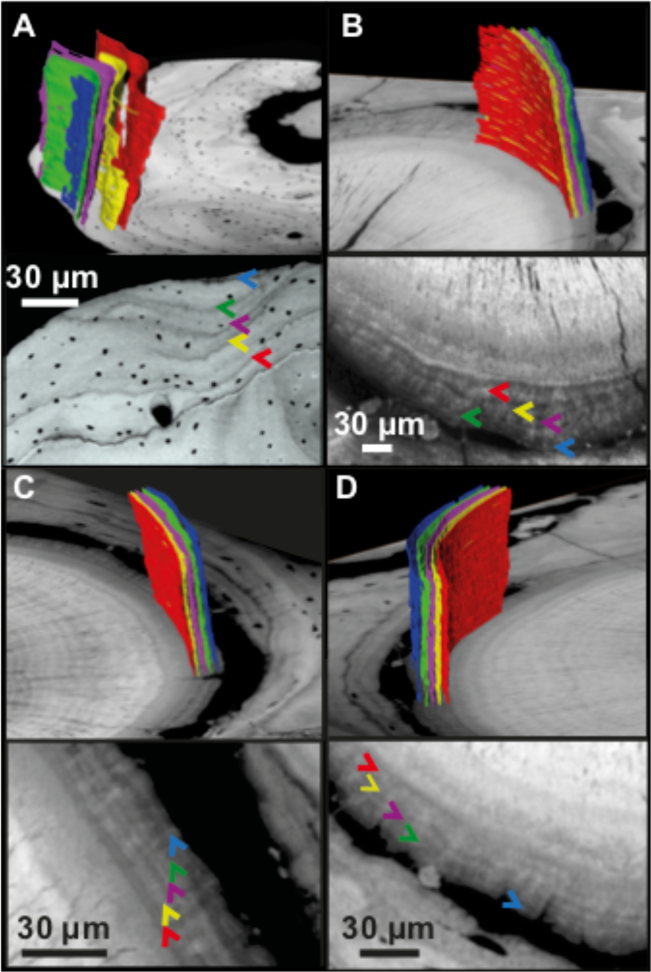
Shared increment patterns between m1 and m2 tooth-root cementum and the dentary of *Morganucodon* specimen NHMUK PV M 96413. **A.** Four lines of arrested growth and a fifth incipient one are visible within the dentary periosteal region, each highlighted by 3D segmented bands of differing colour corresponding to coloured arrows in accompanying transverse PPC-SRµCT slice. Only LAGs persisting through the volume are segmented/highlighted. This pattern is mirrored in **B,** the anterior root of the m1 tooth; **C**, the posterior root of the same m1; and **D,** the anterior root of the m2 tooth.

### Long mammaliaform lifespans and low BMR

Cementum increment counts provide a minimum estimate of maximum lifespan of 14 years for *Morganucodon*, and nine years for *Kuehneotherium* (Figs 2, 4A). These may underestimate true maximum lifespan, as any damage to outer cementum increments would reduce estimated maximum lifespan. One-way ANOVA comparisons of mean intra-observer coefficient of variation (*CV*) between our study and ten previous cementochronological studies of different extant mammal species (Table S3) with similar age ranges show values for PPC-SRµCT data of *Morganucodon* (*CV* = 9.32) and *Kuehneotherium* (*CV* = 4.89) are significantly lower (*p* < 0.01) than previous thin section-based studies (minimum extant *CV =* 14.2, mean *CV* = 21.8).

**Fig. 4.**
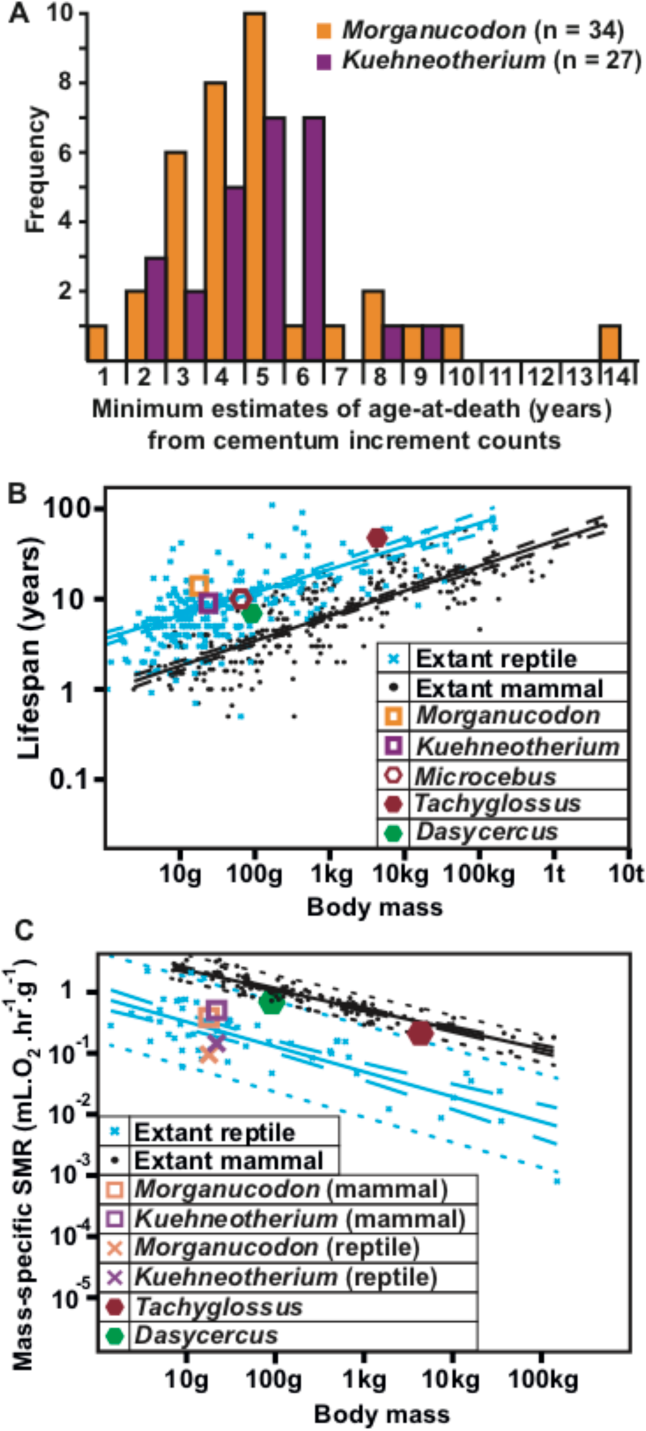
Lifespan and metabolic estimates of *Morganucodon* and *Kuehneotherium.* **A**. Histogram of lifespan estimates from cementum increment counts. **B**. Log_10_ biplot of mean body mass (g) against maximum wild lifespan (years) for extant mammals (n = 279), extant non-avian reptiles (n = 252), and fossil mammaliaforms. **C.** Log_10_ biplot of mean body mass (g) against mass specific standard metabolic rate (msSMR; mL.O_2_.hr^-1^.g^-1^) for extant mammals (n = 117) and extant reptiles (n = 55), and estimates for fossil mammaliaforms. OLS regression lines in **B**, **C,** show extant mammals (black) and extant reptiles (blue), 95% confidence intervals shown with dashed lines, 95% predictor intervals by dotted lines.

We estimated a body mass of 17.9 g for *Morganucodon* and 23.8 g for *Kuehneotherium* (Materials and Methods). Maximum lifespan and mean body mass for the mammaliaforms were compared with published data for large samples of terrestrial, non-volant wild extant mammal (n = 279) and non-avian reptile (n = 252) species (Table S4). Ordinary least squares (OLS) regression of log_10_ transformed values shows the fossil mammaliaforms fall within the range of extant reptiles and have longer maximum lifespans for their size, and are further above the mammal regression mean, than all extant mammals under 4 kg (the long-lived and secondarily dwarfed *(37)* mouse lemur *Microcebus murinus* is closest). Only the short-beaked echidna *Tachyglossus aculeatus*, a monotreme with long lifespan and low metabolic rate, exceeds the *Kuehneotherium*, but not *Morganucodon*, distance above the mammalian mean (Fig 4B). One-way ANCOVA comparisons show that regression slopes for extant mammals and reptiles are statistically similar but their means are significantly separated (*p* <0.01), with reptiles on average living 12.2 years longer than mammals of the same body mass.

To estimate BMR, we used OLS and recovered significant correlations between log10 transformed values of wild lifespan and mass-specific standard metabolic rate (msSMR; measured in mL.O2.hr-1.g-1 and analogous with BMR in extant mammals – SMR was used as BMR cannot be measured in reptiles *(38)*) from published data for 117 extant mammals and 55 extant reptiles (Table S1; Fig S4A). Using the correlation between mammal lifespan and msSMR we estimated a msSMR of 0.38 mL.O2.hr-1.g-1 for *Morganucodon* and 0.51 mL.O2.hr-1.g-1 for *Kuehneotherium.* From the correlation between reptile lifespan and msSMR we estimated a msSMR of 0.10 mL.O2.hr-1.g-1 (*Morganucodon*) and 0.15 mL.O2.hr-1.g-1 (*Kuehneotherium*). When log10 OLS is used to regress these estimates against body mass, both mammaliaforms fall outside the 95% predictor interval (PI) of the mammalian data and within the reptile range of msSMR, whether estimated from mammalian or reptilian data (Fig 4C). This suggests the mammaliaforms had significantly lower msSMR values compared to extant mammals of similar size. The comparably sized mammal (<100 g) of lowest msSMR is the marsupial *Dasycercus cristicauda*, with wild lifespan seven years and msSMR 0.63 mL.O2.hr-1.g-1 (Fig 4C).

In summary, our estimates of maximum lifespan provided by tomographic imaging of cementum increments in *Morganucodon* and *Kuehneotherium* are significantly longer than the wild lifespan of any extant mammal of comparable body mass. These lifespans provide SMR/BMR estimates considerably lower than comparably sized extant mammals, instead corresponding to those of extant reptiles.

### Femoral blood-flow shows intermediate MMR

To compare our fossil mammaliaform BMR estimates with MMR, we used a second proxy directly linked to MMR *(39)*. The ratio between nutrient foramen area and femur length has been used as an index for relative blood flow (*Qi*) through the femur during and after metabolically demanding exercise (*Qi* = *rf*4/*L*; where *rf* = foramen radius and *L* = femur length), previously shown to correlate well with MMR *(39)*. From micro-computed tomography (µCT) data of the six most complete *Morganucodon* femoral diaphyses available, we segmented all nutrient foramina (Fig 5A) and estimated their area by measuring their minimal radii (Materials and Methods). *Kuehneotherium* could not be included as no suitable femoral specimens are known.

**Fig. 5.**
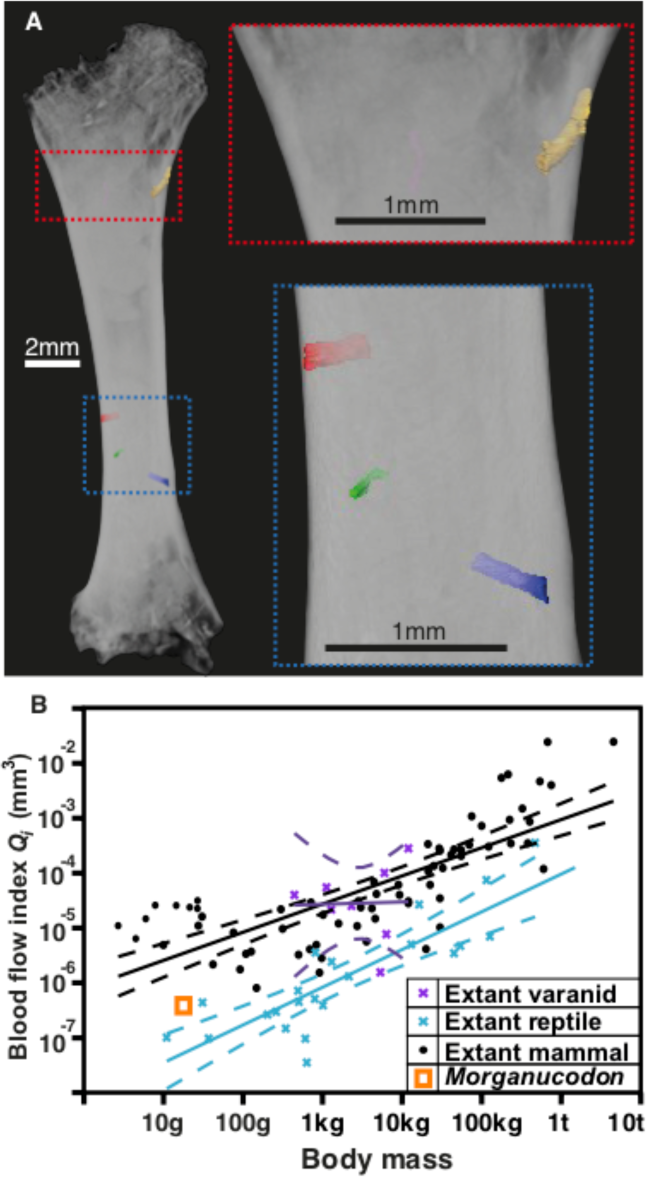
Femoral foramina and estimates of relative femoral bloodflow for *Morganucodon.* **A**. 3D µCT reconstruction of *Morganucodon* femur (specimen UMZC Eo PC 19_6, voxel size 4 μm), with all identifiable foramina segmented/highlighted with colour. **B**. log_10_ biplot of mean body mass (g) against estimated blood flow index (*Qi*; mm^3^) for extant non-varanid reptiles (n = 22), extant varanid reptiles (n=8), extant mammals (n=69), and *Morganucodon*. OLS regression lines in **B** are extant mammals (black), extant non-varanid reptiles (blue) and extant varanids (purple), with 95% confidence intervals represented by dashed lines.

We estimated a *Qi* of 3.829e-7 mm3 for *Morganucodon* and compared this with published and new data (Materials and Methods) for extant mammals (n = 69) and reptiles (n = 30). The latter includes varanids (n = 8) which in the absence of mammalian predators fill an active hunting niche and tend to have mammalian MMR levels while retaining reptilian BMR levels *(39)* (Table S4). One-way ANCOVA comparisons show that means of log10 OLS regression slopes of body mass against *Qi* for extant mammals and non-varanid reptiles are significantly different (*p* <0.01), while the slopes are similar (Materials and Methods). *Morganucodon* is further above (higher *Qi* for its mass) the non-varanid reptile mean than all non-varanid reptiles. However, *Morganucodon* is also slightly further from the mammalian mean than the non-varanid reptile mean, and considerably closer to small non-varanid reptile species than small mammalian species (Fig 5B). This intermediate *Qi*, and so inferred intermediate MMR, suggests that while retaining typical reptilian BMR, *Morganucodon* had an MMR above non-varanid reptiles, but not as high as mammals or actively foraging varanid reptiles.

## Discussion

We have used two quantitative proxies to determine the metabolic status of early mammaliaforms. Relatively long lifespans for both *Morganucodon* and *Kuehneotherium* result in BMR estimates equivalent to modern reptiles of comparable size, and indeed at the higher lifespan/lower BMR end of the reptile scale for *Morganucodon*. In contrast, femoral blood flow estimates (*Q_i_*) suggest that the MMR of *Morganucodon* was intermediate between extant non-varanid reptiles and mammals. We therefore infer that in *Morganucodon* increased MMR (and so also absolute aerobic capacity, AAC = MMR - BMR) was initially selected for before BMR, and that the MMR-first hypothesis *(7)* is the best-supported model for the evolution of mammalian endothermy. We suggest that at least *Morganucodon*, if not also *Kuehneotherium*, occupied a metabolic grade approaching extant varanids; able to undergo longer bouts of aerobically demanding activity than non-varanid reptiles, but not capable of sustaining either mammalian levels of aerobic activity or the elevated thermometabolism exhibited by living endotherms.

Evidence from non-mammalian synapsids (including changes in gait *(8)*, long bone histology *(15)*, and development of secondary osteological features correlated with increased metabolic rate *(18, 19)*) indicate unquestionable changes in physiology from pelycosaur- to mammaliaform-grade taxa. Determinate growth *(40)* and reduction of dental replacement (diphyodonty) in basal mammaliaforms permitted more precise occlusion, which has been considered a key innovation in the development of mammalian endothermy through enabling increased assimilation and higher metabolism *(41)*. However, determinate growth and diphyodonty appear to have preceded the appearance of modern mammalian levels of endothermy, at least in *Morganucodon* and *Kuehneotherium*. We therefore suggest that the development of precise occlusion in basal mammaliaforms may be more associated with dietary specialization and niche partitioning *(1)*.

Comparison of our results to other recent studies of physiology in fossil synapsids supports the hypothesis of a complex, mosaic pattern of endothermic evolution with different characters being selected for at different rates through time and with respect to phylogeny. For example, the size diminution associated with the cynodont-mammaliaform transition *(42)* may have reversed the evolutionary trajectory of some previous histological proxies for endothermy *(43)*, contributing to the complex, contradictory patterns observed. Our study also suggests more work is needed to directly compare fossil and extant ecto- and endothermic taxa, to better understand their relative metabolic properties. Many previous studies rely on simple binary divisions, such as the presence/absence of fibrolamellar bone and/or respiratory nasal turbinates. These proxies cannot represent accurately the complex series of physiological characteristics that range between ectothermy and endothermy, and are frequently distributed homoplastically across the synapsid phylogeny, individually and with respect to each other. Other studies with relative data such as preserved apatite oxygen isotopes *(22)* allow comparisons with co-habiting ectothermic taxa but cannot be directly compared to extant data and cannot indicate where fossil taxa fall in the metabolic spectrum of extant vertebrates. However, our results are compatible with recent work on living mammals that the BMR of the Middle Jurassic (∼170 Ma) mammalian MRCA was comparable to present-day values *(14)*. This indicates evolution towards modern-day mammalian endothermy occurred during the 25 million year-long Early Jurassic and suggests the mammalian mid-Jurassic adaptive radiation *(5, 6)* was driven by this, or vice versa.

In conclusion, our data offer a direct link to measurable aspects of endothermy, BMR and MMR, at a key point in mammalian evolution. Further work applying these methods to younger Mesozoic mammaliaforms and mammals, and comparison with evidence from other physiological characteristics, will allow the evolutionary tempo and mode of multiple aspects of mammalian physiology to be determined. The early mammaliaforms *Morganucodon* and *Kuehneotherium* possessed surprisingly low, reptile-like metabolic rates, plus a mixture of plesiomorphic and derived characters *(7)* relating to life history and physiology. Ultimately, we can no longer assume that the endothermic metabolism of living mammals had evolved in the earliest mammaliaforms.

## Acknowledgements

Many thanks to the Natural History Museum, London, University Museum of Zoology, Cambridge, and the Finnish Museum of Natural History, Helsinki, Finland, for loans of specimens; facilitated by Martha Richter, Rob Asher and Matt Lowe, and Martti Hildén. We thank Keijo Hämäläinen for his help in the initial stages of the synchrotron imaging. Assistance with lab work and materials: Tom Davies, Wendy Dirks, Liz Martin-Silverstone, Dani Schmidt, Remmert Schouten, Pedro Viegas, John Cunningham & Duncan Murdock. Discussions: Chris Dean, Wendy Dirks, Jim Hopson, Thomas Martin, Rachel O’Meara, Stephen Naji, Tanya Smith & Emily Rayfield.

## Author Contributions

IJC & PGG conceived and designed the project. EN, PGG, PB, KR & IJC selected, prepared and curated specimens. EN, PGG, PB, VF, DH, TK, AK, FM, AP, BP, PS, HS, PT & IJC performed the synchrotron experiments. EN, AK & IJC performed the microCT experiments. EN processed and EN, CN & KY analysed the synchrotron data. EN & IJC analysed the micro-CT data. EN, PGG & IJC discussed the interpretations. EN wrote the first draft and created all figures; EN, PGG & IJC wrote the manuscript; all authors provided a critical review of the manuscript and approved the final draft. This article originated as a Masters thesis (University of Bristol), then a Ph.D. thesis (University of Southampton), performed by EN and supported and supervised by PGG, PS, NJG, JJ, KRB, MJB & IJC. Authors MJB to KW contributed equally to this work and are listed alphabetically.

## Data and materials availability

The tomographic data that support the findings of this study are available from the corresponding authors upon reasonable request. Physiological source data for charts and graphs in figures 4 and 5b are provided with the paper in Supplementary Materials tables.

## Supplementary Materials

Materials and Methods

Supplementary text

Figures S1-S5

Tables S1-S5

References (*44-321*)

