## Supplement for "Reptile-like physiology in Early Jurassic stem-mammals" (from another version of this preprint: version 1, 2019-09-30)

**This PDF file includes:**

Materials and Methods
Supplementary Text
Figs. S1 to S5
Tables S1 to S5

### Materials and Methods

**Choice of fossil specimens.** All *Morganucodon* and *Kuehneotherium* specimens are from the Early Jurassic St Brides Island fissure suite, from Glamorgan, South Wales, UK *(1, 44)*. The two genera co-existed on a small landmass during the Early Jurassic marine transgression (Hettangian-early Sinemurian, ~200 Ma) *(45)*. Thousands of their bones and teeth were washed into karst fissures that have subsequently been revealed by quarrying. The *Morganucodon* and *Kuehneotherium* specimens used for cementum analysis are from the Natural History Museum, London, UK (NHMUK) and *Morganucodon* femora specimens from the University Museum of Zoology, Cambridge, UK (UMZC). *Morganucodon* specimens are *Morganucodon watsoni* from Pontalun 3 fissure in Pontalun quarry (now Lithalun), excavated in 1962-1963 *(44, 45)*. *Kuehneotherium* specimens are from three Glamorgan fissures in Pontalun and Pant quarries; Pontalun 3, Pant 2 and Pant 4. Although the Pant 2 *Kuehneotherium* collection is more extensive, teeth are poorly mineralized and so produced relatively fewer usable increment counts. Pontalun 3 and Pant 2 fissures have a relatively impoverished fauna, and Pant 4 has a more diverse biota *(45)*. *Kuehneotherium* from Pontalun 3 fissure is *Kuehneotherium praecursoris (46),* but those from Pant fissures are considered different species *(47)*. Pontalun 3 fissure specimens in NHMUK were prepared by immersing dried blocks of clay matrix in hot water, adding dilute hydrogen peroxide or sodium hexametaphosphate (Calgon) if required, but those in UMZC were prepared with 10% acetic acid. *Kuehneotherium* specimens from Pant quarry (Pant 2 and Pant 4) were collected by the University College London Kermack team, from 1955-1978, and are from harder matrix, prepared with 15% acetic acid *(47)*.

*Morganucodon* and *Kuehneotherium* specimens used for cementum analysis are either isolated teeth, or dentary specimens with a range of teeth *in-situ.* The lower dental formula of

*Morganucodon* is i4: c1: p4; m4-5 and for *Kuehneotherium* i?: c1: p6: m6 *(1)*. In *Morganucodon,* most isolated teeth measured were lower second molars (m2) since they are easily identified, relatively large, have robust separated roots, and, as anterior diphyodont molars, should erupt relatively early and are not replaced, therefore offering a near-complete record of life history. However, i4, c, p3, p4, m1, m3, and m4 teeth were also studied in dentulous *Morganucodon* specimens. *Kuehneotherium* is not abundant from Pontalun 3 fissure, and so specimens were also selected from two Pant quarry fissures. Dentulous *Kuehneotherium* specimens are extremely rare, so mainly isolated teeth were used. There are relatively fewer *Kuehneotherium* mid row lower molars with well-preserved cementum than for *Morganucodon*, so all appropriate teeth were chosen; p5, p6, and a range of lower and upper molars *(47)*.

**Tomographic imaging of cementum**. Pilot scans of two *Morganucodon* lower second molars (NHMUK PV M 104131 and NHMUK PV M 104132) were carried out on the nanotomography imaging beamline ID22 at the European Synchrotron Radiation Facility (ESRF), Grenoble, France (project EC 1064). For ID22 we used these experimental settings for computed tomographic (CT) imaging: X-ray energy of 29.6 keV, 1999 projections over 180° rotation, 0.5 ms exposure time, 321 nm voxel size, 405 mm sample-to-detector distance, diamond window, a 20 µm thick LSO scintillator doped with Tb, and 1.5 mm Al filter, single propagation distance tomography.

71 additional *Morganucodon* specimens (52 isolated teeth and 19 dentaries) and two pilot *Kuehneotherium* specimens were scanned at the ID19 beamline of ESRF (project ES 152). A single harmonic U13 undulator X-ray source was used, delivering a pink X-ray beam with peak energy at 26.5 keV, with 1.4 mm Al filter used to cut background lower energies. The detector was a microscope optic system coupled to a sCMOS sensor (PCO edge 5.5), mounted with a 10

μm thick GGG:$_{Eu}$ scintillator. Scans were performed using single propagation distance tomography (15 mm sample-to-detector propagation distance), exposure time of 250-300 ms, 2499 angular projections over 360°, and at voxel sizes of 280, 347 and 700 nm.

117 additional *Kuehneotherium* specimens (116 isolated teeth and one dentary) and 12 additional *Morganucodon* dentary specimens were scanned at the TOMCAT tomographic beamline of the Swiss Light Source (SLS), Villigen, Switzerland. The beam was set at 20 keV using a double multilayer monochromator, a LSO:Tb scintillator and a pco.EDGE 5.5 detector. Samples were scanned using single propagation distance tomography (14 mm sample-to-detector propagation distance), exposure time 150 ms, and 1500 angular projections over $180^0$ at voxel size 330 nm. A *Kuehneotherium* lower molar (UMZC Sy 141) was imaged at 1.2 μm voxel size (with exposure time 150 ms and 1500 angular projections over 180°) to provide the 3D volume in Fig 1B.

Three juvenile *Morganucodon* dentary specimens with roots from final deciduous premolars (NHMUK PV M 27312, NHMUK PV M 27474 and NHMUK PV M 27475), and an older individual with extensive molar wear (NHMUK PV M 27465), were scanned during a separate experiment at TOMCAT beamline, SLS. Energy was 21 keV using a double multi-layer monochromator. Samples were scanned using single propagation distance tomography (14 mm sample-to-detector propagation distance), exposure time 200 ms and 1601 angular projections over 180° at voxel size 330 nm.

CT reconstructions of resulting tomographic data were generated using a filtered back-projection algorithm coupled with 'Paganin-style' single distance phase retrieval *(48)* algorithms developed in-house at the respective beamlines *(49)*. For data from ID19, $\beta = 8.1\ 10^{-8}$, $\delta = 9.8\ ^{10-9}$. For data from TOMCAT, $\beta = 3.7\ 10^{-8}$, $\delta = 1.7\ 10^{-10}$.

Of 71 *Morganucodon* specimens imaged at beamline ID19, four were additionally imaged at the ESRF nano-imaging beamline ID16A (project ES 152). These were imaged using holotomography *(50)* from four propagation distances. Holograms were recorded using a CCD detector with a physical pixel size of 1.5 μm and a 23 μm thick GGG:Eu scintillator at both 15 keV and 33.6 keV. Voxel sizes were 10 nm, 25 nm, 30 nm and 130 nm. Angular projections recorded over $180^0$ varied between 1200 and 2000, exposure times were 250 to 800 ms. To generate the image containing the virtual section in Fig 1G, with 30 nm voxel size, the four focus-to-sample distances were 2.65 mm, 13.19 mm, 15.36 mm and 19.87 mm and sample-to-detector distance was 1.2648 m.

For the 3D model in Fig 1A, a *Morganucodon* lower molar (NHMUK PV M 104134) was imaged using micro CT (μCT) at the University of Helsinki using a Nanotom 180 NF (phoenix|X-ray Systems & Services GmbH) with a CMOS detector (Hamamatsu Photonics) and a high-power transmission-type X-ray nanofocus source with tungsten anode. 900 angular projections were collected over 180°, with exposure time of 1840 ms and voxel size 2 μm. Raw projection data were reconstructed using filtered back-projection by datos|x rec reconstruction software, supplied by the system manufacturer.

**Increment counting and the creation of virtual thin sections**. Cementum increments were counted in CT data using modifications to techniques suggested by the Cementochronology Research Program[32] to account for the 3D nature of the PPC-SRμCT cementum data. First, cementum was visually inspected throughout the volume of each scan, in transverse PPC-SRμCT slices using ImageJ/Fiji *(51)* to distinguish between specimens that were confidently interpreted as preserving cementum increments, and those too badly affected by diagenesis for increment counting. Phase contrast imaging of incremental features is understood to be prone to recurrent destructive interference patterns from Fresnel diffraction that create periodic blurring at differing frequencies when they are scanned using inappropriate experimental parameters (principally X-ray energy, sample-to-detector propagation distance and voxel size). However, our parameters produce blurring frequencies that are too narrow (approx. 500-900 nm) to significantly affect contrast between cementum increments (1-3 μm radial thickness). Therefore, no significant masking of increments from Fresnel diffraction blurring should be expected in our data. In specimens that preserved increments, volumes were inspected by eye to identify regions of highest increment contrast with no lensing and/or coalescence between increments. Increments identified in these regions were followed by eye throughout the entire cementum tissue surrounding these regions, both longitudinally and transversely through the root, to distinguish between principal increments and accessory increments formed by lensing and coalescence (Fig S1). Principal increments were distinguished as those that persisted vertically through the entire scanned region of cementum, whereas accessory increments lasted only for short periods before coalescing into neighbouring increments (Fig S1).

Once regions of highly contrasting primary cementum increments were identified, virtual thin sections of these were created by isolating 10 transverse PPC-SRμCT slices through each region and summing their greyscale values using the "Sum slices" option of the "Z projection" tool in ImageJ/Fiji. This created a new image of increased contrast between dark and light cementum increments, and reduced image noise. Three-to-five virtual thin sections were created for all specimens with readable cementum increments. For each virtual thin section, increments were counted manually by three observers: Observer One (EN) had considerable experience in counting cementum increments (>100 specimens studied); Observer Two (KW) had training in counting cementum increments (30 specimens studied under guidance from Observer One) and experience in studying growth patterns in PPC-SRμCT data of long-bones; Observer Three (CN) had no prior experience in counting increments or studying growth patterns. Each observer studied virtual thin sections blind, following numbering and randomization between specimens using the RAND function in Microsoft Excel. For each observer, final specimen increment counts were determined as the maximum number counted in all their virtual thin sections.

Precision between observers increment counts (Fig S5, Table S1) was compared by calculating the coefficient of variation (*CV*; formula 1) (Table S3):

1. $$CV = \left[\frac{standard\ deviation}{mean}\right] * 100$$

**3D modelling of cementum increments**. 3D modelling was performed, using Avizo image analysis software (Avizo 8.0; Thermo Fisher Scientific), on a subsample of teeth: the first tooth imaged using PPC-SRμCT (NHMUK PV M 104131), specimens with the highest cementum increment count for each fossil taxon (NHMUK M 104127 for *Morganucodon* and UMCZ Sy 141 for *Kuehneotherium*), and a *Morganucodon* specimen with both dentary lags and cementum increments (NHMUK PV M 96413). Original tomographic data was downsampled in each axis by a factor of two to decrease manual processing time while retaining sufficient spatial resolution to preserve cementum increments. Principal increments originally identified by eye were manually traced in each PPC-SRμCT slice and assigned to different materials in the 'label field' tool-kit. Models were created and analysed using the 'surface view' feature of Avizo. This allowed the pattern of incrementation to be viewed in 3D, to test whether increments defined in 2D slices were true principal increments or diffuse accessory increments (Fig S1).

**Body mass estimation for *Morganucodon* and *Kuehneotherium* and choice of extant taxa for physiological comparison.** **B**ody mass of our fossil taxa were estimated using two techniques, based on scaling between single cranial dimensions and measured body mass in extant mammals. Maximum body mass estimates were made using the scaling relationship between dentary length (mm) and body mass (g) published for extant marsupial mammals of small body mass, and subsequently used to estimate body mass in several Mesozoic fossil taxa from the Late Jurassic Morrison Formation and elsewhere *(52)*:

**2.** $$\ln \text{body mass(g)} = 2.9677*(\ln \text{dentary length(mm)}) - 5.6712$$

Dentary lengths are from published CT reconstructions *(1)* (20 mm for *Morganucodon* and 21.9 mm for *Kuehneotherium*), resulting in body mass estimates of 25.0 g and 32.7 g respectively. However, this estimate may be an overestimate for *Kuehneotherium* as it has a longer, more gracile dentary relative to *Morganucodon*, due to their differing feeding ecologies *(1)*.

Secondly, minimum body mass estimates were calculated using the scaling relationship between skull length (mm) and body mass (g) found for 64 extant species of small 'lipotyphlan' insectivores *(53)* and used to estimate the body masses of several Mesozoic mammaliaforms *(20, 53, 54)*.

**3.** $$\text{Ln(body mass(g))} = 3.68*(\ln(\text{skull length(mm)}) - 3.83$$

Due to a lack of complete, diagnostic cranial material for UK samples of *Morganucodon* and *Kuehneotherium*, skull lengths were estimated using a scaling relationship between dentary length and skull length calculated for *Morganucodon oehleri* from Fig S1 *(19)* (skull length(mm) = 1.0458*jaw length(mm)). Following this relationship, we used dentary length to estimate a skull length of 21.0 mm for *Morganucodon watsoni* and 22.9 mm for *Kuehneotherium*, resulting in body mass estimates of 10.7 g and 14.9 g respectively (as above this may be an overestimate for *Kuehneotherium*). Body masses presented in Fig S3 and S4 are mean values of the two estimates (17.9 g for *Morganucodon*, 23.8 g for *Kuehneotherium*).

Information on maximum wild lifespan and mean body mass was obtained for 278 extant terrestrial mammal species (body mass: mean = 70.2 kg, S.D. = 395 kg; wild lifespan: mean = 10.8 years, S.D. = 11.2 years) and 256 extant terrestrial non-avian reptile species (body mass: mean = 3.6 kg, S.D. = 20 kg; wild lifespan: mean = 12 years, S.D. = 14 years) (SI Table 1). Information for flying or gliding taxa (including birds) was not included as their body masses are known to be secondarily reduced to aid their lifestyle and so may distort observed overall relationships between lifespan and body mass *(55-57)*; birds and bats live on average three-to-four times as long as terrestrial mammals of similar mass *(58)*. Marine mammals were not included as their environment positively affects available body mass, allowing significantly higher body masses than terrestrial taxa *(59-61)*. Mammal wild lifespan data is from primary literature; the majority from a download from (https://www.demogr.mpg.de/longevityrecords/0203.htm) on 10/03/2019. Most reptile lifespans come from the supplementary information of *(62)*. Body mass estimates were from an online *Ecological Archives* database *(63)*. (http://www.esapubs.org/archive/ecol/E084/094/metadata.htm), the Anage database (https://genomics.senescence.info/species/), and published data *(38, 62)*.

Data on standard metabolic rate (SMR) for extant mammals (n=117; mean = 915; S.D. = 3018) and reptiles (n=55; mean = 62.3; S.D. = 159) was obtained from a published electronic appendix *(38)*. SMRs for mammals were included only if measured under basal metabolic conditions *(38)* and so are synonymous with basal metabolic rates.

Data on the post-natal growth rate constant *K* was obtained upon request for extant mammals *(64)* (n = 115, mean = 0.03, S.D. = 0.03), and reptiles (n=33, mean = 0.002, S.D. = 0.003) from previous publications *(65, 66)*. Growth rate is measured in days$^{-1}$, and the dimensionless constant *K* is considered the best measure for comparing growth rates between multiple species *(67)*.

**µCT study of femoral nutrient foramina.** Femoral nutrient foramina in *Morganucodon* and 11 extant mammal taxa of comparable size were µCT imaged in the X-Ray Micro-Imaging Laboratory, University of Helsinki (Table S4). We used µCT data of femoral foramina rather than photographic images of foramina *(39)* because many of the *Morganucodon* foramina were sediment-filled, making photographic measurement difficult/impossible, whereas the differential density of sediment and fossilized bone in µCT data allowed visualization and analysis. All femora (fossil and extant) were scanned using a Bruker Skyscan 1272 µCT scanner at 70 kVp source tension and with 0.5mm Al filter. 1125-1800 angular projections were collected over 180° with exposure time 1344 ms. One fossil specimen was scanned at isotropic voxel size 4 µm (Fig 4A), and five specimens at 5 µm. All femora from extant taxa were scanned at 4µm voxel sizes. Raw projection data were reconstructed using filtered back-projection with the Feldkamp algorithm by the Bruker reconstruction software "NRecon" (Version 1.7.1.0).

Analysis of µCT femora data was conducted using Avizo image analysis software (Version 9.3.0; Thermo Fisher Scientific). Foramina were located using the "Orthoslice" tool to scroll through transverse CT slices. Once located, foramina were imaged in 3D using the "Volume rendering" tool, allowing the minimum diameter, and so minimum radius (in cm), of the foramen aperture to be measured using the "3D line measurement" tool. Minimum diameter was used *(39)*, in the study of foramina in small mammals, as it is not always possible to see the direction of nutrient vessel penetration. Resulting foramina radii were used to generate estimates of blood flow index ($Q_i$) following the method outlined in *(39)*. If multiple foramina were found in a single specimen, their radii were summed following the assumption that this represents the total entry/exit potential of nutrient circulation through the femur *(39)*. Single or summed radii (abbreviated as $r$; measured in cm), were used with femur length ($L$; measured in cm) to estimate $Q_i$:

**4.** $$\boldsymbol{Q_i = r^4/L}$$

Although *Morganucodon* femoral elements are relatively common in the *Hirmeriella* fissure suite, almost all are incomplete *(68)*. We used the six most complete femoral specimens (UMZC EoPC 19_1 to EoPC 19_6) from the Pontalun 3 fissure where *Morganucodon* teeth imaged using PPC-SRµCT originated, which preserve at least two thirds of the femoral shaft. Length was estimated from the minimum mid-shaft diaphysial width using the scaling relationship between these established for a reconstructed *Morganucodon* femur created by concatenating three incomplete femora:

**5. Femur length = 10.3 * maximum mid-diaphysial diameter**

**Statistics**. One-way ANOVA comparison of intra-observer CV between cementochronological studies; Shapiro-Wilks normality test: PPC-SRµCT data $W = 1$, $p = 1$, histological data $W = 0.93$, $p = 0.41$; test statistics F = 11.12, degrees of freedom(df) = 10, Cohen's effect size $d = 3.13$, $p = 0.00728$. One-way ANCOVA comparison of OLS regression slopes for lifespan against body mass in mammals ($\log_{10}$ lifespan = 0.274($\log_{10}$ body mass) - 0.05; 95% CI = 0.255, 0.292; $r^2 = 0.69$) and reptiles ($\log_{10}$ lifespan = 0.253($\log_{10}$ body mass) + 0.465; 95% CI = 0.225, 0.281; $r^2 = 0.46$); slopes are statistically similar ($F = 1.09$, $p = 0.3$) while means are significantly separated ($F = 243$, df = 529, partial eta squared effect size = 0.32, $p = 2.3E^{-45}$). OLS regression of mammalian lifespan against msSMR: $\log_{10}$ msSMR = -0.623($\log_{10}$ lifespan) + 0.298; 95% CI = -0.71, -0.526; $r^2 = 0.59$, $p = 0.0001$). OLS regression of reptilian lifespan against msSMR: $\log_{10}$ msSMR = -0.494($\log_{10}$ lifespan) – 0.067; 95% CI = -1.328, -0.479; $r^2 = 0.43$, $p = 0.0001$). OLS regression of mammalian lifespan against growth constant $K$; $\log_{10} K$ = -0.99($\log_{10}$ lifespan) – 0.972; 95% CI = -1.138, -0.853; $r^2 = 0.66$, $p$ <0.0001). OLS regression of reptilian lifespan against growth constant $K$; $\log_{10} K$ = -0.948($\log_{10}$ lifespan) – 1.937; 95% CI = -1.4, -0.415; $r^2 = 0.43$, $p = 0.0002$). One-way ANCOVA comparison of OLS regression slopes for $Q_i$ against body mass in extant mammals ($\log_{10}(Qi)$ = 0.513*$\log_{10}$(body mass) - 6.104) and non-varanid reptiles ($\log_{10}(Qi)$ = 0.685*$\log_{10}$(body mass) - 8.139); slopes are statistically similar ($F = 2$, $p = 0.16$) while means are significantly different ($F = 87.6$, df = 89, partial eta squared effect size = 0.50, $p = 7.4E^{-15}$).

### Supplementary Text

#### Taxonomic description of *Morganucodon* and *Kuehneotherium*

Order MORGANUCODONTA Kermack, Mussett and Rigney, 1973

Family MORGANUCODONTIDAE Kühne 1958

Genus and species MORGANUCODON WATSONI Kühne, 1949

Family KUEHNEOTHERIIDAE Kermack, Kermack and Mussett, 1968

Genus and species KUEHNEOTHERIUM PRAECURSORIS Kermack, Kermack and Mussett, 1968

The phylogenetic relationship of *Morganucodon* and *Kuehneotherium* is shown in Fig S2. The position of *Morganucodon* at the base of Mammaliaformes (Node 4) has remained stable between multiple studies *(1, 19, 28, 69)*. We have also chosen to position *Kuehneotherium* at the Mammaliaformes node in Fig S2, as its lack of diagnostic post-cranial material makes its phylogenetic placement uncertain and poorly supported amongst studies *(28, 47, 69)*. However, it is generally placed either within a clade including *Morganucodon (47. 69)*, or further crownward than *Morganucodon* based mainly based on the triangulation of its molars *(28, 47, 69)*.

**Eruption sequence and timing**

As noted in the main text, establishing individual ages from several teeth along *Morganucodon* lower jaws provides information on the tooth eruption sequence and timing. Eight scanned dentaries provided information (Table S2). There are several specimens with tooth row increment counts covering p1 to m2: For example, NHMUK PV M 95790 for p1 to p3; NHMUK PV M96413 for p3 to m1; NHMUK PV M 96396 for p4 to m2 (Table S2). NHMUK PV M 96413 and NHMUK PV M 96396 also show readable lines of arrested growth (LAGs) in the dentary bone, which provide the same count as their p3 to m1 and p4-m2 teeth respectively. This indicates that p1 to m2 erupted within one year of each other in *Morganucodon*.

There is also a specimen (NHMUK PV M 27312) with an increment count of one (Table S1). This specimen has the deciduous fourth premolar in situ, with signs of the developing permanent third and fourth premolars forming, but also the first molar *in situ* and m2 in the process of erupting. This shows that m1 erupted before p3 and 4, and all within one year of life. In the mammaliaform *Docodon victor* the third premolar p3 is the earliest tooth position for premolar replacement *(70)*, but that level of differentiation cannot yet be determined in *Morganucodon*.

Three specimens (NHMUK PV M 96396; NHMUK PV M 104129; NHMUK PV M 104130) show that the third molar erupted during the year after the permanent p1-m2 erupted. One specimen (NHMUK PV M 95790) has cementum increment counts indicating that the permanent final incisor i4, and the canine, erupted during the year after the eruption of the first three permanent premolars. We therefore conclude that the ultimate incisor, canine and the third molar erupted during the year following the eruption of p1 to m2 in *Morganucodon*. We do not have information on eruption timing of more anterior incisors, or the fourth molar. The fifth molar is only very occasionally present.

The eruption of the final incisor and canine after the premolars may be a plesiomorphic feature in *Morganucodon* and does not follow the general antero - posterior pattern seen in later therians. In the Late Jurassic dryolestids *(71)*, tooth replacement takes place in two waves; with i2, i4, p1, and p3 in the first series and i1, i3, c, p2, and p4 in the second. However, this is also different from the pattern seen here in *Morganucodon*, where the ultimate incisor and canine come in after the permanent premolars.

*Morganucodon* could live to at least 14 years, commonly to six and not unusually to nine years, so this pattern of most of the adult tooth row being in place during the first two years of life is also supportive of a relatively short (compared with its total lifespan) juvenile stage *(69)* and determinate growth as established by O'Meara and Asher *(40)*. The absolute length of these stages in *Morganucodon* is however considerably longer than extant mammals of comparable body size *(31)*.

Unfortunately, there are no available tooth rows of *Kuehneotherium* with cementum increment counts. Dentulous specimens are rare and the three specimens in the UMZC collection are not suitable for scanning for cementum annuli as they were previously mounted in plaster and dissected to reveal the tooth roots. *Kuehneotherium* has a longer tooth row than *Morganucodon* with six premolars and six molars *(47).* A difference from *Morganucodon* is that there is evidence from specimens of juvenile *Kuehneotherium* dentaries *(47),* that the canine, ultimate premolar, p6, and third molar all erupted at about the same time, which suggests that the canine erupted relatively earlier in *Kuehneotherium* than in *Morganucodon*.

The loss of the anterior permanent postcanines has been noted in *Morganucodon* jaws *(72)*, and is assumed to proceed along the tooth row with age. In some dentaries, the first premolar appears to be lost relatively early and is only represented by a small, indistinct, resorbed root just posterior to the canine. Accordingly, it is not certain if the timing, and degree, of loss of the anterior premolars is variable in individuals, but this question is part of a current study. Specifically, the specimen NHMUK PV M 95790 with cementum annuli counts for i4 to p3, obviously has not shed the anterior premolars, in spite of a minimum age of eight years.

**Using lifespan to estimate growth rates in stem mammals**

We investigated a final third proxy for metabolic potential in fossil mammaliforms; their post-natal growth rate. Growth rates have been shown to correlate strongly with metabolic power in extant vertebrates, with endotherms growing an order of magnitude faster than ectotherms (*30, 65)*, and show significant correlation with lifespan in terrestrial vertebrates *(64-67).* We estimated growth rates using Ordinary Least Squares correlations between wild lifespan and growth rate from published data from 115 extant mammals *(64)* and 31 extant reptiles *(65, 66)* (Fig. S4B). From mammal data we estimate growth rate constants *K* ($days^{-1}$ - Methods) of $7.68e^{-3}$ $days^{-1}$ (*Morganucodon*) and $1.19e^{-2}$ $days^{-1}$ (*Kuehneotherium*). From reptile data we estimated $K$ = $9.53e^{-4}$ $days^{-1}$ (*Morganucodon*) and $K = 1.44e^{-3}$ $days^{-1}$ (*Kuehneotherium*). $Log_{10}$ OLS regression against body mass again places both mammaliaforms outside the mammalian 95% PI and within the reptile growth rate range, whether estimated from mammalian or reptilian data (Fig. S4C). The lowest growth rate of any <100g extant mammal is $K = 3.24e^{-2}$ $days^{-1}$ for the Mongolian gerbil *Meriones unguiculatus* (Fig. S4C).

Fig. S1.

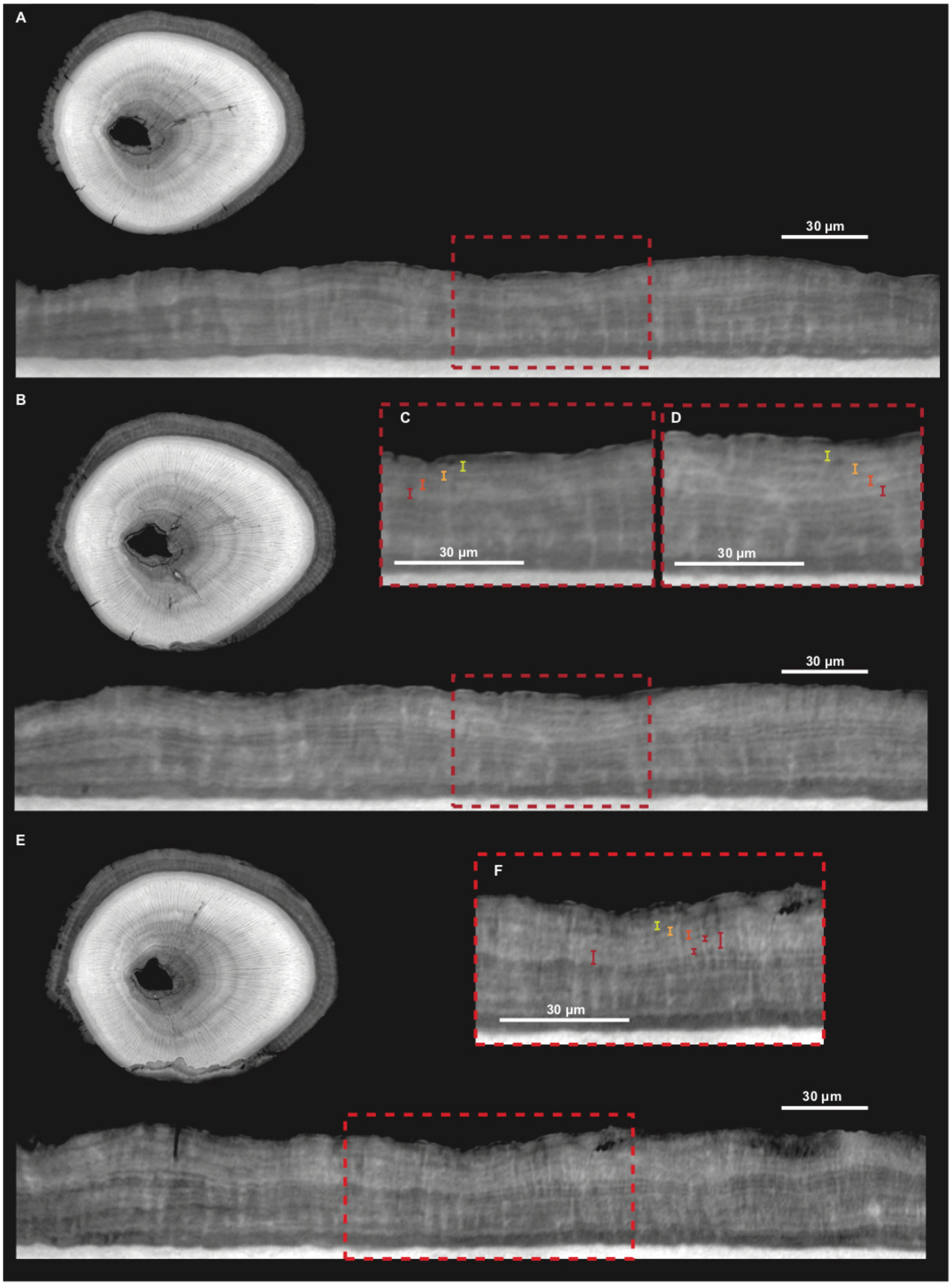

**Fig. S1. Example of lensing and coalescence of cementum increments in specimen NHMUK PV M 104138. A**. Transverse PPC-SRµCT slice selected from region of the cementum closest to the crown. Inset to the left is the entire slice, and the straightened portion represents the region of highest increment contrast from this slice. **B.** Transverse PPC-SRµCT slice 100 µm towards the root apex relative to **A**. **C.** Detail from **A** highlighted by dashed red box, with four outermost higher density (lighter coloured) cementum increments annotated with coloured bracketed lines. **D**. Detail from **B** highlighted by dashed red box. The same increments imaged in **A** and **C** are annotated with the same coloured bracketed lines. However, they are more clearly defined in **B** and **D**, with the innermost two annotated increments **(**dark red and orange bracketed lines) coalescing in **A** and **C**. **E.** Transverse PPC-SRµCT slice 100 µm towards the root apex relative to **B. F**. Detail from **E** highlighted by dashed red box. While the four outermost increments imaged in the other PPC-SRµCT slices are also imaged here, the innermost increment (dark red bracketed lines) has lensed, creating two accessory increments. All straightening performed using the "straighten" tool in ImageJ/Fiji *(51)*.

**Fig. S2.**

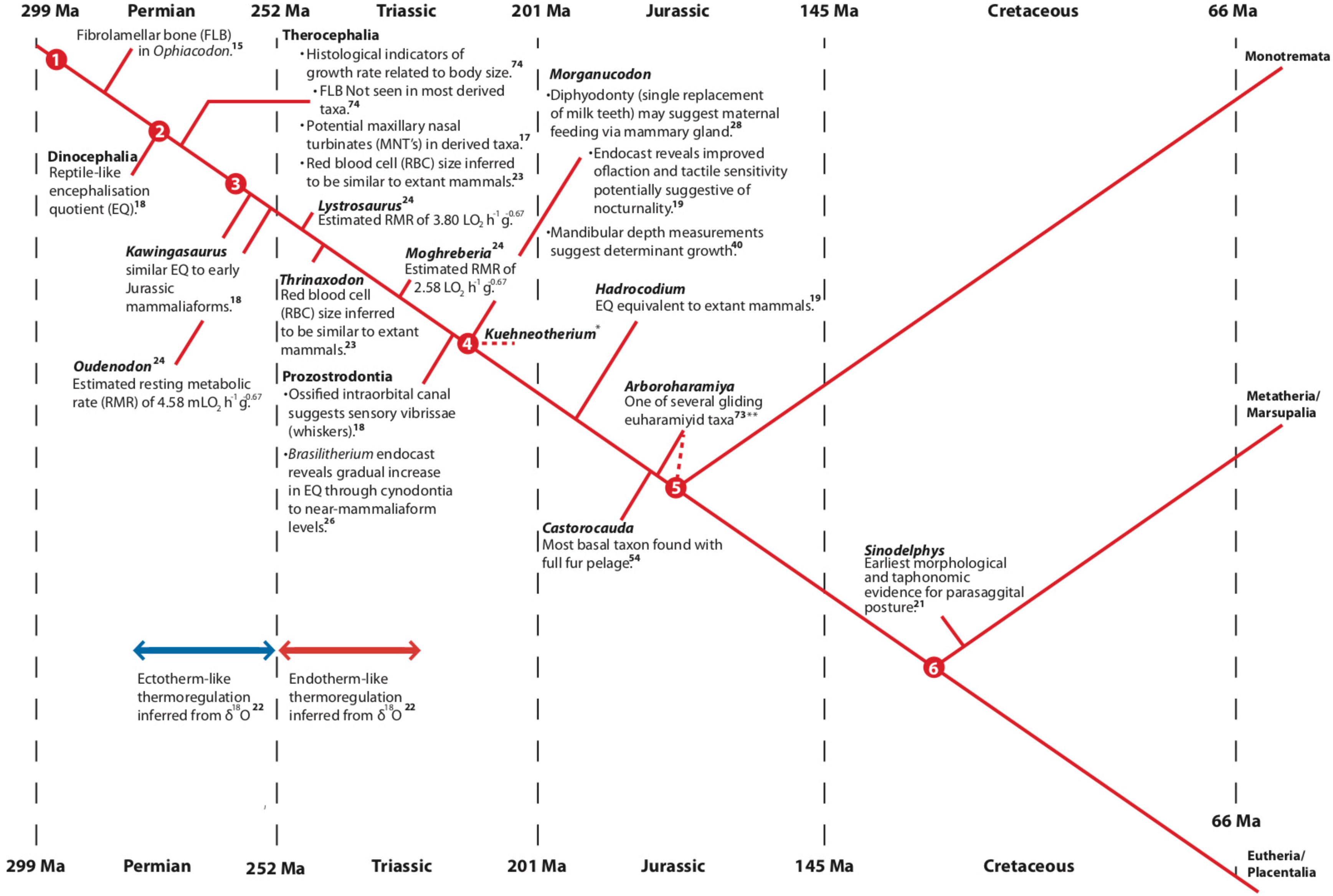

299 Ma
Permian
252 Ma
Triassic
201 Ma
Jurassic
145 Ma
Cretaceous
66 Ma
Fibrolamellar bone (FLB) in Ophiacodon.15
Dinocephalia
Reptile-like encephalisation quotient (EQ).18
Therocephalia
•Histological indicators of growth rate related to body size.74
•FLB Not seen in most derived taxa.74
•Potential maxillary nasal turbinates (MNT's) in derived taxa.17
•Red blood cell (RBC) size inferred to be similar to extant mammals.23
Lystrosaurus24
Estimated RMR of 3.80 LO2 h-1 g-0.67
Kawingasaurus
similar EQ to early Jurassic mammaliaforms.18
Oudenodon24
Estimated resting metabolic rate (RMR) of 4.58 mLO2 h-1 g-0.67
Thrinaxodon
Red blood cell (RBC) size inferred to be similar to extant mammals.23
Moghreberia24
Estimated RMR of 2.58 LO2 h-1 g-0.67
Prozostrodontia
•Ossified intraorbital canal suggests sensory vibrissae (whiskers).18
•Brasilitherium endocast reveals gradual increase in EQ through cynodontia to near-mammaliaform levels.26
Morganucodon
•Diphyodonty (single replacement of milk teeth) may suggest maternal feeding via mammary gland.28
•Endocast reveals improved oflaction and tactile sensitivity potentially suggestive of nocturnality.19
•Mandibular depth measurements suggest determinant growth.40
Kuehneotherium*
Hadrocodium
EQ equivalent to extant mammals.19
Arboroharamiya
One of several gliding euharamiyid taxa73**
Castorocauda
Most basal taxon found with full fur pelage.54
Sinodelphys
Earliest morphological and taphonomic evidence for parasaggital posture.21
Monotremata
Metatheria/ Marsupalia
Eutheria/ Placentalia
Ectotherm-like thermoregulation inferred from δ18O 22
Endotherm-like thermoregulation inferred from δ18O 22
1
2
3
4
5
6

**Fig. S2. Timescaled phylogeny summarising evidence for physiological evolution along the synapsid lineage towards mammals.** Red nodes highlight the divergence of major lineages; Node 1 = divergence of the Pelycosaur lineage; Node 2 = the Therapsida clade; Node 3 = the Cynodontia clade; Node 4 = the Mammaliaformes clade; Node 5 = the Mammalia clade; Node 6 = the Theria clade. Superscript numbers denote references in the main text. * Denotes the uncertain phylogenetic affinities of *Kuehneotherium* within the Mammaliaformes clade *(28).* ** Denotes the uncertain phylogenetic affinities of *Arboroharamiya (73).*

**Fig. S3.**

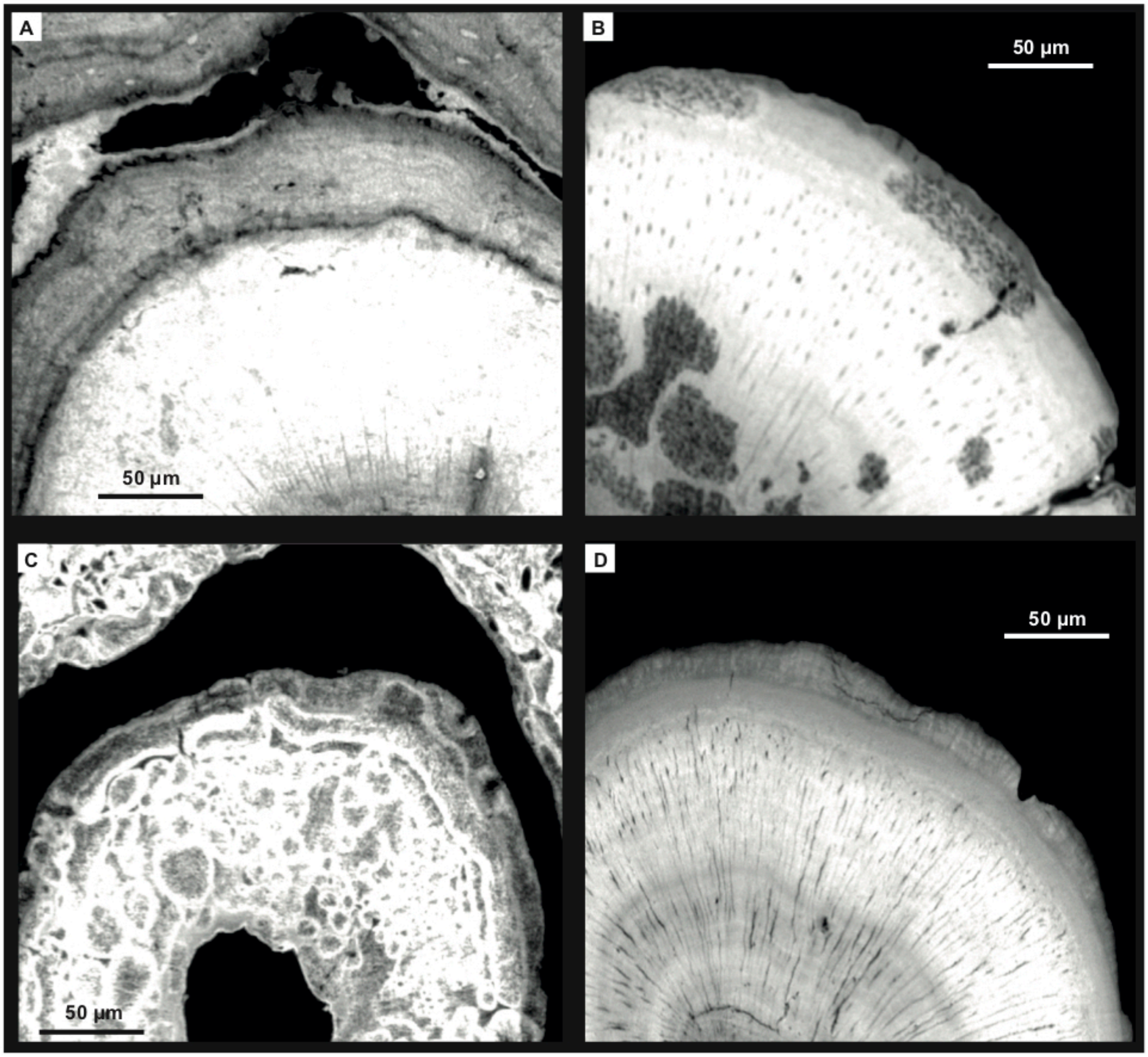

**Fig. S3. Common biological and physical features, and diagenetic fabrics, encountered in tomographic data of fossil cementum. A.** Substantial variation in the thickness of individual cementum increments in the anterior root of the m2 specimen NHMUK PV M 104129. **B.** Discrete dark, less dense regions of diagenetic alteration within the root of NHMUK PV M 96086, a specimen of otherwise excellent dentine and cementum preservation. **C.** Globular diagenetic fabrics have adulterated virtually all microstructure in the anterior root of the m1 specimen NHMUK PV M 95809, though it may still be possible to separate dentine and cementum. **D**. Physical damage to the cementum tissue has removed outer increments in discrete regions of the cementum of the anterior root of NHMUK PV M 96273. The dentine has been over-saturated (white) by decreasing the dynamic range in imageJ/Fiji *(51)*, to improve the visibility of the cementum.

**Fig. S4.**

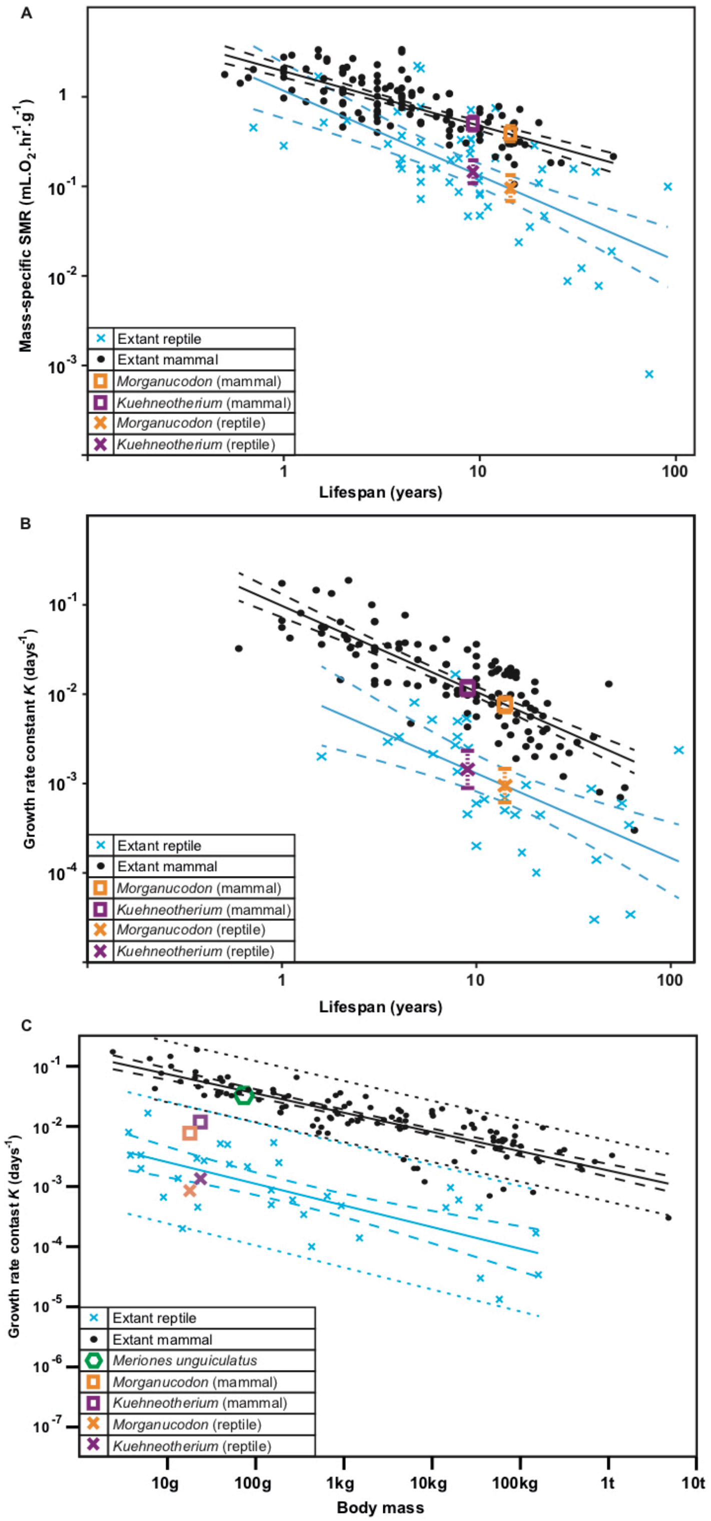

A
Mass-specific SMR (mL.O$_2$.hr$^{-1}$.g$^{-1}$)
Lifespan (years)
Extant reptile
Extant mammal
*Morganucodon* (mammal)
*Kuehneotherium* (mammal)
*Morganucodon* (reptile)
*Kuehneotherium* (reptile)
B
Growth rate constant *K* (days$^{-1}$)
Lifespan (years)
Extant reptile
Extant mammal
*Morganucodon* (mammal)
*Kuehneotherium* (mammal)
*Morganucodon* (reptile)
*Kuehneotherium* (reptile)
C
Growth rate contast *K* (days$^{-1}$)
Body mass
10g
100g
1kg
10kg
100kg
1t
10t
Extant reptile
Extant mammal
*Meriones unguiculatus*
*Morganucodon* (mammal)
*Kuehneotherium* (mammal)
*Morganucodon* (reptile)
*Kuehneotherium* (reptile)

**Fig. S4. The relationship between lifespan and msSMR, and lifespan and growth rate constant *K*, in mammals and reptiles. A.** $Log_{10}$ biplot between maximum wild lifespan (years) and mass-specific standard metabolic rate (msSMR; $mL.O_2.hr^{-1}.kg^1$) for extant mammals (n=117) and reptiles (n=55). **B.** $Log_{10}$ biplot between maximum wild lifespan (years) and post-natal growth rate constant *K* ($days^{-1}$) for extant mammals (n=115) and reptiles (n=31). **C.** $Log_{10}$ biplot of mean body mass (g) against post-natal growth rate constant *K* ($days^{-1}$) for extant mammals (n = 115) and extant reptiles (n = 33) and estimates for fossil mammaliaforms. OLS regression means for each clade (black lines for mammals, blue lines for reptiles) are used to estimate msSMR (**A**), and *K* (**B**) for mammaliaforms *Morganucodon* and *Kuehneotherium*. 95% confidence intervals are represented by dashed lines, 95% predictor intervals by dotted lines in **C**.

**Fig. S5.**

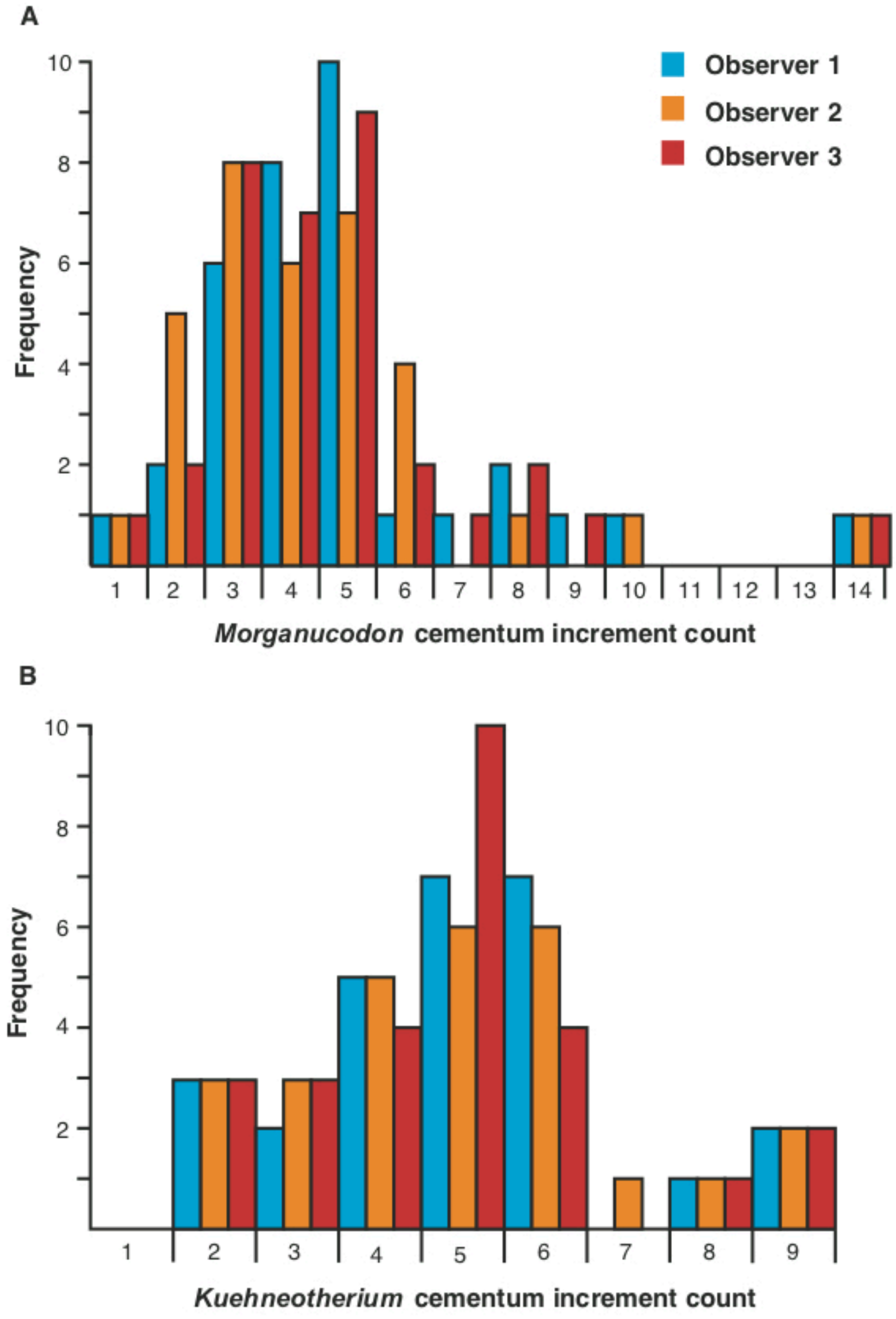

**Fig. S5. Comparison between distributions of cementum increment counts estimated by three independent observers. A.** Distributions of increment counts estimated for *Morganucodon.* **B.** Distributions of increment counts estimated for *Kuehneotherium.*

### Table S1

| Taxon | Specimen | Observer 1 increment count | Observer 2 increment count | Observer 3 increment count |
|---|---|---|---|---|
| *Morganucodon* | NHMUK PV M 104126 | 5 | 6 | 5 |
| *Morganucodon* | NHMUK PV M 104127 | 14 | 14 | 14 |
| *Morganucodon* | NHMUK PV M 104128 | 5 | 5 | 5 |
| *Morganucodon* | NHMUK PV M 104129 | 9 | 5 | 6 |
| *Morganucodon* | NHMUK PV M 104130 | 5 | 5 | 5 |
| *Morganucodon* | NHMUK PV M 104131 | 4 | 3 | 3 |
| *Morganucodon* | NHMUK PV M 104132 | 4 | 3 | 4 |
| *Morganucodon* | NHMUK PV M 104133 | 3 | 3 | 3 |
| *Morganucodon* | NHMUK PV M 104134 | 4 | 4 | 4 |
| *Morganucodon* | NHMUK PV M 104135 | 5 | 5 | 5 |
| *Morganucodon* | NHMUK PV M 104136 | 2 | 2 | 2 |
| *Morganucodon* | NHMUK PV M 104137 | 5 | 3 | 4 |
| *Morganucodon* | NHMUK PV M 104138 | 10 | 10 | 9 |
| *Morganucodon* | NHMUK PV M 27312 | 1 | 1 | 1 |
| *Morganucodon* | NHMUK PV M 92528 | 3 | 2 | 3 |
| *Morganucodon* | NHMUK PV M 95790 | 8 | 8 | 8 |
| *Morganucodon* | NHMUK PV M 95805 | 8 | 4 | 8 |
| *Morganucodon* | NHMUK PV M 95809 | 3 | 2 | 3 |
| *Morganucodon* | NHMUK PV M 95810 | 5 | 6 | 5 |
| *Morganucodon* | NHMUK PV M 96075 | 8 | 8 | 8 |
| *Morganucodon* | NHMUK PV M 96085 | 3 | 2 | 3 |
| *Morganucodon* | NHMUK PV M 96086 | 4 | 3 | 4 |
| *Morganucodon* | NHMUK PV M 96141 | 5 | 5 | 3 |
| *Morganucodon* | NHMUK PV M 96380 | 4 | 4 | 4 |
| *Morganucodon* | NHMUK PV M 96396 | 4 | 4 | 4 |
| *Morganucodon* | NHMUK PV M 96406 | 2 | 2 | 2 |
| *Morganucodon* | NHMUK PV M 96413 | 5 | 5 | 5 |
| *Morganucodon* | NHMUK PV M 96418 | 7 | 6 | 7 |
| *Morganucodon* | NHMUK PV M 96441 | 5 | 3 | 5 |
| *Morganucodon* | NHMUK PV M 96444 | 3 | 3 | 3 |
| *Morganucodon* | NHMUK PV M 96463 | 6 | 6 | 6 |
| *Morganucodon* | NHMUK PV M 96464 | 4 | 4 | 4 |
| *Morganucodon* | NHMUK PV M 96474 | 4 | 4 | 5 |
| *Morganucodon* | NHMUK PV M 96476 | 5 | 5 | 5 |
| *Morganucodon* | NHMUK PV M 96490 | 3 | 3 | 3 |
| *Kuehneotherium* | NHMUK PV M 20982 | 5 | 6 | 5 |
| *Kuehneotherium* | NHMUK PV M 20990 | 3 | 3 | 3 |
| *Kuehneotherium* | NHMUK PV M 21069 | 6 | 6 | 6 |
| *Kuehneotherium* | NHMUK PV M 21080 | 5 | 5 | 4 |
| *Kuehneotherium* | NHMUK PV M 21081 | 2 | 2 | 2 |
| *Kuehneotherium* | NHMUK PV M 27257 | 4 | 3 | 4 |
| *Kuehneotherium* | NHMUK PV M 27273 | 5 | 4 | 5 |
| *Kuehneotherium* | NHMUK PV M 27301 | 2 | 2 | 2 |
| *Kuehneotherium* | NHMUK PV M 27305 | 2 | 2 | 2 |
| *Kuehneotherium* | NHMUK PV M 27308 | 6 | 6 | 5 |
| *Kuehneotherium* | NHMUK PV M 27344 | 6 | 5 | 5 |
| *Kuehneotherium* | NHMUK PV M 27436 | 4 | 4 | 4 |
| *Kuehneotherium* | NHMUK PV M 27443 | 4 | 4 | 4 |
| *Kuehneotherium* | NHMUK PV M 27505 | 6 | 6 | 5 |
| *Kuehneotherium* | NHMUK PV M 27520 | 5 | 5 | 5 |
| *Kuehneotherium* | NHMUK PV M 27529 | 5 | 5 | 5 |
| *Kuehneotherium* | NHMUK PV M 27537 | 5 | 5 | 5 |
| *Kuehneotherium* | NHMUK PV M 27538 | 6 | 6 | 6 |
| *Kuehneotherium* | NHMUK PV M 27545 | 6 | 6 | 6 |
| *Kuehneotherium* | NHMUK PV M 27550 | 4 | 5 | 5 |
| *Kuehneotherium* | NHMUK PV M 27628 | 5 | 4 | 5 |
| *Kuehneotherium* | NHMUK PV M 27653 | 9 | 9 | 9 |
| *Kuehneotherium* | NHMUK PV M 27745 | 6 | 7 | 6 |
| *Kuehneotherium* | NHMUK PV M 27816 | 3 | 3 | 3 |
| *Kuehneotherium* | UCMZ Sy 106 | 8 | 8 | 8 |
| *Kuehneotherium* | UCMZ Sy 121 | 4 | 4 | 3 |
| *Kuehneotherium* | UCMZ Sy 141 | 9 | 9 | 9 |

**Table S1.** Independent estimates of cementum increment counts estimated by three observers.

Observer 1 = EN; observer 2 = CN; observer 3 = KW.

**Table S2**

| Specimen | Element | Observer 1 count |
|---|---|---|
| NHMUK PV M 95790 | i4 | 7 |
| NHMUK PV M 95790 | c | 7 |
| NHMUK PV M 95790 | p1 | 8 |
| NHMUK PV M 95790 | p2 | 8 |
| NHMUK PV M 95790 | p3 | 8 |
| NHMUK PV M 96413 | p3 | 5 |
| NHMUK PV M 96413 | p4 | 5 |
| NHMUK PV M 96413 | m1 | 5 |
| NHMUK PV M 96413 | dentary | 5 |
| NHMUK PV M 96396 | p4 | 4 |
| NHMUK PV M 96396 | m1 | 4 |
| NHMUK PV M 96396 | m2 | 4 |
| NHMUK PV M 96396 | m3 | 3 |
| NHMUK PV M 96396 | dentary | 4 |
| NHMUK PV M 95809 | m1 | 3 |
| NHMUK PV M 95809 | m2 | 3 |
| NHMUK PV M 104128 | m1 | 5 |
| NHMUK PV M 104128 | m2 | 5 |
| NHMUK PV M 96441 | m1 | 5 |
| NHMUK PV M 96441 | m2 | 5 |
| NHMUK PV M 104130 | m1 | 5 |
| NHMUK PV M 104130 | m2 | 5 |
| NHMUK PV M 104130 | m3 | 4 |
| NHMUK PV M 104129 | m1 | 9 |
| NHMUK PV M 104129 | m2 | 9 |
| NHMUK PV M 104129 | m3 | 8 |

**Table S2.** Cementum and dentary increment counts for each element of dentulous *Morganucodon* specimens. The lower dental formula of *Morganucodon* is i4: c: p4: m4-5.

**Table S3**

| Study | Taxon | *n* | Maximum age (years) | mean *CV* |
|---|---|---|---|---|
| Newham et al. (presented here) | *Morganucodon* | 34 | 14 | 9.32 |
| Newham et al. (presented here) | *Kuehneotherium* | 27 | 9 | 4.89 |
| Grau et al.[75] | *Procyon lotor* | 54 | 9 | 20.6 |
| Gasawey et al.[76] | *Alces alces* | 72 | 9 | 14.2 |
| Klevezal and Pucek[77] | *Bison bonasus* | 45 | 21 | 20.57 |
| Kay and Cant[78] | *Macaca mulatta* | 65 | 24 | 29.34 |
| Cederlund et al.[79] | *Capreolus capreolus* | 74 | 9 | 30.5 |
| Bodkin et al.[80] | *Enhydra lutris* | 14 | 14 | 26.24 |
| Landon et al.[81] | *Canis lupus* | 12 | 6.8 | 25.92 |
| Christensen-Dalsgaard et al.[82] | *Ursus maritimus* | 32 | 15 | 15.2 |
| Pasda[83] | *Rangifer tarandus* | 63 | 16 | 19.4 |
| Perez-Barberia et al.[84] | *Cervus elaphus* | 164 | 17 | 16.22 |

**Table S3.** Results of quantitative analyses of precision between three independent observers for the cementum increment counts of fossil mammals studied here, *Morganucodon* and *Kuehneotherium*, and those of ten previous studies of extant mammals with comparable age ranges. *CV* – coefficient of variation. Superscript numbers denote references in the main text.

**Table S4**

| Taxon | Average femur length (cm) | Average foramen radius (cm) | Average $Q_i$ ($mm^3$) |
|---|---|---|---|
| *Apodemus flavicolis* | 2.15 | 0.082 | 3.15E-05 |
| *Micromys minutus* | 1.1 | 0.062 | 1.47E-05 |
| *Microtus levis* | 1.43 | 0.071 | 2.26E-05 |
| *Mus musculatus* | 1.32 | 0.073 | 2.38E-05 |
| *Peromyscus truei* | 1.75 | 0.063 | 1.10E-05 |
| *Sorex minutissimus* | 0.48 | 0.045 | 1.00E-05 |
| *Sicista betulina* | 1.04 | 0.071 | 2.59E-05 |
| *Sorex araneus* | 0.82 | 0.044 | 4.89E-06 |
| *Myodes rutilus* | 1.17 | 0.066 | 1.61E-05 |
| *Neomys fodiens* | 1.04 | 0.068 | 2.28E-05 |
| *Sorex minutus* | 0.56 | 0.042 | 6.37E-06 |
| *Morganucodon* | 1.25 | 0.026 | 3.83E-07 |

**Table S4.** Results of measuring femur length and nutrient foramen radius in μCT data for 11 extant mammal taxa and *Morganucodon*, with resulting estimates of blood flow index $Q_i$.

**Table S5**

| Order | Taxon | Aka | Body mass (g) | Body mass ref. | Alt. body mass (g) | Alt. body mass ref. | Alt. body mass (g) | Alt. body mass ref. | Wild lifespan (years) | SMR ($mL.O_2.hr^{-1}$) | Growth rate constant ($day^{-1}$) | Wild lifespan ref. |
|---|---|---|---|---|---|---|---|---|---|---|---|---|
| Mammalia | *Acinonyx jubatus* | Cheetah | 37900 | 38 | | | | | 12 | 8099.751 | | 219 |
| Mammalia | *Acomys cahirinus* | Spiny mouse | 42 | 38 | 45.2 | 64 | | | 1.6 | 48.6511 | 0.05623 | 273 |
| Mammalia | *Acomys russatus* | Golden spiny mouse | 55.4 | 86 | | | 55.55 | 38 | 3 | 46.35892 | | 273 |
| Mammalia | *Aepyceros melampus* | Impala | 52500 | 64 | | | | | 9 | | 0.0043 | 296 |
| Mammalia | *Ailuropoda melanoleceuca* | Giant panda | 125000 | 86 | | | | | 26 | | | 115 |
| Mammalia | *Ailurus fulgens* | Red panda | 4325 | 64 | 5740 | 85 | | | 14 | 915.3 | 0.009 | 168 |
| Mammalia | *Alcelaphus buselaphus* | Hartebeest | 159000 | 86 | | | | | 20 | | | 211 |
| Mammalia | *Alces alces* | Moose | 325000 | 86 | | | | | 27 | | | 259 |
| Mammalia | *Alouatta palliata* | Mantled howler | 4670 | 86 | | | | | 20 | | | 163 |
| Mammalia | *Alouatta seniculus* | Red howler | 6543 | 86 | | | | | 25 | | | 271 |
| Mammalia | *Antechinomys laniger* | Kultarr | 25.8 | 38 | | | | | 3.3 | 31.90611 | | 316 |
| Mammalia | *Antechinus flavipes* | Yellow-footed antechinus | 46.5 | 38 | | | | | 3.5 | 61.50761 | 0.0351 | 291 |
| Mammalia | *Antechinus stuartii* | Brown antechinus | 25 | 86 | | | 28.2 | 38 | 2 | 45.75579 | | 118 |

| | | | | | | | | | | | | |
|---|---|---|---|---|---|---|---|---|---|---|---|---|
| Mammalia | *Antechinus swainsoni* | Dusky antechinus | 66.9 | 38 | | | | | 2 | 77.27176 | | 147 |
| Mammalia | *Antidorcas marsupialis* | Springbok | 39000 | 64 | | | | | 20 | | 0.0137 | 85 |
| Mammalia | *Antilocapra americana* | Pronghorn | 46100 | 64 | | | 34779 | 85 | 10 | | 0.0083 | 219 |
| Mammalia | *Antilope cervicapra* | Blackbuck | 37500 | 86 | | | | | 18 | | | 267 |
| Mammalia | *Aotus trivirgatus* | Night/owl monkey | 820 | 38 | | | | | 20 | 422 | | 271 |
| Mammalia | *Aplodontia rufa* | Mountain beaver | 630 | 38 | | | | | 10 | 277.2 | | 219 |
| Mammalia | *Apodemus agrarius* | Striped field mouse | 21.2 | 86 | | | | | 1.5 | | | 188 |
| Mammalia | *Apodemus sylvaticus* | Old World wood and field mouse | 23.9 | 38 | | | | | 1 | 49.48212 | | 196 |
| Mammalia | *Arctocebus calabarensis* | Golden potto | 206 | 86 | | | | | 9.5 | | | 183 |
| Mammalia | *Arvicola amphibius* | European water vole | 94.7 | 86 | | | | | 2 | | | 274 |
| Mammalia | *Atelerix frontalis* | African hedgehog | 555 | 86 | | | | | 4 | | | 185 |
| Mammalia | *Baiomys taylori* | Northern pygmy mouse | 7.2 | 86 | 8 | 64 | 7.15 | 38 | 1.1 | 21.01403 | 0.0425 | 155 |
| Mammalia | *Bassariscus astutus* | Ringtail | 1015 | 64 | | | | | 7 | | 0.0215 | 85 |
| Mammalia | *Bettongia penicillata* | Woylie | 1018 | 38 | | | | | 6 | 609.8379 | | 129 |

| | | | | | | | | | | | | |
|---|---|---|---|---|---|---|---|---|---|---|---|---|
| Mammalia | *Bison bonasus* | European bison | 610000 | 86 | | | | | 20 | | | 204 |
| Mammalia | *Blarina brevicauda* | Northern short-tailed shrew | 21.6 | 64 | 20.9 | 85 | | | 2.2 | | 0.188 | 85 |
| Mammalia | *Burramys parvus* | Mountain pygmy possum | 44.3 | 38 | | | | | 4 | 44.75014 | | 245 |
| Mammalia | *Callimico goeldii* | Goeldi's monkey | 555 | 64 | | | | | 9 | | 0.0097 | 182 |
| Mammalia | *Callithrix jacchus* | Common marmoset | 190 | 85 | 255.2 | 64 | | | 12 | | 0.0172 | 182 |
| Mammalia | *Canis latrans* | Coyote | 10000 | 38 | 13250 | 64 | | | 14.5 | 2979.196 | 0.0183 | 107 |
| Mammalia | *Canis lupus* | Wolf | 26625 | 64 | | | | | 16 | | 0.0177 | 201 |
| Mammalia | *Canis mesomelas* | Black-backed jackal | 10250 | 64 | | | | | 10 | 3860 | 0.0149 | 216 |
| Mammalia | *Capra ibex* | ibex | 82500 | 64 | | | | | 16 | | 0.005 | 108 |
| Mammalia | *Castor fiber* | Eurasian beaver | 25,000 | 85 | | | | | 24 | | | 164 |
| Mammalia | *Cavia porcellus* | Guinea pig | 629 | 38 | | | | | 14.8 | 312.1045 | | 117 |
| Mammalia | *Ceratotherium simum* | White rhinoceros | 2E+06 | 85 | | | | | 50 | | | 175 |
| Mammalia | *Cercartetus nanus* | Eastern pygmy possum | 64.8 | 85 | | | 70 | 38 | 5 | 77.154 | | 312 |
| Mammalia | *Cercocebus albigena* | Gray-cheeked mangabey | 9318 | 85 | | | | | 21 | | | 184 |

| | | | | | | | | | | | | |
|---|---|---|---|---|---|---|---|---|---|---|---|---|
| Mammalia | *Cercocebus atys* | Sooty mangabey | 8600 | 86 | | | | | 18 | | | 184 |
| Mammalia | *Cercocebus torquatus* | Collared mangabey | 9493 | 85 | | | | | 20.5 | | | 184 |
| Mammalia | *Cercopithecus diana* | Diana monkey | 4550 | 85 | | | | | 34.8 | | | 184 |
| Mammalia | *Cervus elaphus* | Red deer | 67000 | 85 | 200000 | 64 | | | 16.1 | | 0.006 | 240 |
| Mammalia | *Chaetodipus formosus* | Long-tailed pocket mouse | 15.1 | 85 | | | | | 4 | | | 207 |
| Mammalia | *Chaetodipus penicillatus* | Desert pocket mouse | 17.35 | 85 | | | | | 2.16 | | | 234 |
| Mammalia | *Chinchilla lanigera* | Chinchilla | 426 | 38 | | | 436.7 | 85 | 11.3 | 253.9436 | | 117 |
| Mammalia | *Choloepus didactylus* | Southern two-toed sloth | 6250 | 64 | | | | | 25 | | 0.002 | 302 |
| Mammalia | *Clethrionomys glareolus* | Bank vole | 20.8 | 64 | 21.5 | 85 | | | 1.25 | | 0.0809 | 307 |
| Mammalia | *Colobus polykomos* | Western black and white colobus | 9525 | 85 | | | | | 30.5 | | | 271 |
| Mammalia | *Connochaetes taurinus* | Blue wildebeest | 164500 | 64 | 196500 | 85 | | | 21 | | 0.0026 | 109 |
| Mammalia | *Cratogeomys castanops* | Mexican pocket gopher | 270 | 85 | | | | | 2.6 | | | 295 |
| Mammalia | *Crocidura leucodon* | Bicolored shrew | 11 | 64 | | | | | 2.9 | | 0.1 | 85 |

| | | | | | | | | | | | | |
|---|---|---|---|---|---|---|---|---|---|---|---|---|
| Mammalia | *Crocidura russula* | Greater white-toothed shrew | 10.8 | 85 | 11.6 | 64 | | | 1.5 | | 0.146 | 222 |
| Mammalia | *Crocuta crocuta* | Spotted hyena | 63000 | 64 | | | | | 19 | | 0.0056 | 149 |
| Mammalia | *Cryptotis parva* | Least shrew | 6.3 | 85 | 5.5 | 64 | | | 1.8 | | 0.134 | 264 |
| Mammalia | *Ctenomys talarum* | Tuco-tuco | 121 | 38 | | | | | 2 | 133.4216 | | 124 |
| Mammalia | *Cynomys ludovicianus* | Black-tailed prairie dog | 1112 | 38 | 1125 | 64 | | | 8 | 483.4789 | 0.0111 | 189 |
| Mammalia | *Damaliscus dorcas* | Sassaby | 84500 | 85 | | | | | 21.4 | | | 196 |
| Mammalia | *Damaliscus pygargus* | Blesbok | 84500 | 64 | | | | | 10 | | 0.0056 | 164 |
| Mammalia | *Dasycercus cristicauda* | Mulgara | 91 | 85 | | | 101 | 38 | 7 | 57.44462 | | 317 |
| Mammalia | *Dasypus novemcinctus* | Common long-nosed armadillo | 3510 | 38 | | | | | 15 | 1242.45 | | 318 |
| Mammalia | *Dasyuroides byrnei* | Kowari | 91.7 | 38 | | | | | 4 | 95.984815 | | 135 |
| Mammalia | *Dasyurus geoffroii* | Western quoll | 1327 | 85 | | | | | 5 | 684.9998 | | 282 |
| Mammalia | *Dasyurus hallucatus* | Northern quoll | 571 | 85 | | | 558 | 38 | 3 | 301.9363 | | 250 |
| Mammalia | *Diceros bicornis* | Black rhinoceros | 1100000 | 64 | | | | | 40 | | 0.0033 | 253 |
| Mammalia | *Didelphis marsupialis* | Southern opossum | 1244 | 85 | | | 1165 | 38 | 2 | 778.1077 | | 300 |

| | | | | | | | | | | | | |
|---|---|---|---|---|---|---|---|---|---|---|---|---|
| Mammalia | *Didelphis virginiana* | Large American opossum | 2847 | 85 | 3000 | 64 | 2488 | 38 | 3 | 1138.385 | 0.0129 | 192 |
| Mammalia | *Dipodomys heermani* | Heermann's kangaroo rat | 63.3 | 85 | | | | | 3 | | | 166 |
| Mammalia | *Dipodomys merriami* | Merriam's kangaroo rat | 37.6 | 85 | 42 | 64 | 36.5 | 38 | 4 | 47.08432 | 0.0368 | 207 |
| Mammalia | *Dipodomys nitratoides* | Fresno kangaroo rat | 37.8 | 85 | | | | | 3 | | | 166 |
| Mammalia | *Dipodomys ordii* | Ord's kangaroo rat | 47.8 | 85 | | | 46.8 | 38 | 2 | 91.25668 | | 178 |
| Mammalia | *Echinops telfairi* | Lesser Madagascar “hedgehog” | 116.4 | 85 | | | | | 1.2 | | | 196 |
| Mammalia | *Elaphurus davidianus* | Pere David's deer | 186500 | 85 | | | | | 23.3 | | | 195 |
| Mammalia | *Elephantulus myurus* | Long-eared elephant shrew | 62.97 | 38 | | | | | 1.1 | 74.60186 | | 292 |
| Mammalia | *Elephantulus rufescens* | Rufous elephant shrew | 53 | 38 | 58 | 64 | | | 1.6 | 61.13734 | 0.0481 | 292 |
| Mammalia | *Elephas maximus* | Asiatic elephant | 4E+06 | 85 | | | | | 70 | | | 208 |
| Mammalia | *Eligmodontia typus* | Highland gerbil mouse | 17.5 | 38 | | | | | 0.7 | 35.30826 | | 170 |

| | | | | | | | | | | | | |
|---|---|---|---|---|---|---|---|---|---|---|---|---|
| Mammalia | *Eliomys quercinus* | Garden dormouse | 82.5 | 64 | | | | | 2.17 | | 0.0409 | 99 |
| Mammalia | *Enhydra lutris* | Sea otter | 26833 | 85 | | | | | 23 | | | 272 |
| Mammalia | *Equus burchellii* | Burchell's zebra | ###### | 64 | | | | | 22 | | 0.0038 | 293 |
| Mammalia | *Equus zebra* | Mountain zebra | ###### | 85 | | | | | 24 | | | 174 |
| Mammalia | *Erethizon dorsatum* | North American porcupine | 8600 | 64 | | | 6871 | 85 | 18 | | 0.0079 | 150 |
| Mammalia | *Erinaceus europaeus* | Eurasian hedgehog | 750 | 38 | | | | | 7 | 510.3632 | | 237 |
| Mammalia | *Erinaceus europaeus* | Western European hedgehog | 1214 | 85 | | | | | 7 | 510.3632 | | 237 |
| Mammalia | *Felis chaus* | Jungle cat | 10000 | 64 | | | | | 14 | | 0.0165 | 251 |
| Mammalia | *Fukomys damarensis* | Damaraland mole-rat | 131.3 | 85 | | | 138 | 38 | 8.5 | | | 278 |
| Mammalia | *Galago senegalensis* | Senegal bushbaby | 192.2 | 64 | 171.7 | 85 | | | 16 | 139.35 | 0.0169 | 182 |
| Mammalia | *Galea musteloides* | Yellow-toothed cavy | 322 | 38 | | | | | 1.3 | 283.8147 | | 271 |
| Mammalia | *Galemys pyrenaicus* | Pyrenean desman | 57.5 | 85 | | | | | 3.5 | | | 269 |
| Mammalia | *Galerella sanguinea* | Slender mongoose | 519.6 | 85 | | | 540 | 38 | 10 | 381.7477 | | 172 |
| Mammalia | *Geocapromys ingrahami* | Bahamian hutia | 775 | 85 | | | | | 8 | | | 228 |

| | | | | | | | | | | | | |
|---|---|---|---|---|---|---|---|---|---|---|---|---|
| Mammalia | *Geomys bursaris* | Plains pocket gopher | 197 | 38 | | | | | 4 | 188.0479 | | 137 |
| Mammalia | *Georychus capensis* | Cape mole-rat | 195 | 38 | | | | | 3 | 136.4734 | | 304 |
| Mammalia | *Gerbillus gerbillus* | Lesser gerbil | 29.7 | 38 | | | | | 2.25 | 46.1332 | | 288 |
| Mammalia | *Gerbillus pyramidum* | Greater Egyptian gerbil | 37 | 64 | 108.5 | 85 | | | 2.25 | 99.04 | 0.0329 | 288 |
| Mammalia | *Giraffa camelopardalis* | Giraffe | 800000 | 85 | | | | | 26 | | | 141 |
| Mammalia | *Glis glis* | Fat dormouse | 174.4 | 85 | | | | | 9 | | | 260 |
| Mammalia | *Gorilla gorilla* | Gorilla | 139842 | 64 | | | | | 43 | | 0.0008 | 120 |
| Mammalia | *Gymnobelideus leadbeateri* | Leadbeater's possum | 166 | 85 | | | | | 7.5 | | | 212 |
| Mammalia | *Herpestes auropunctatus* | Javanese mongoose | 611 | 85 | | | | | 6.75 | 336.1698 | | 97 |
| Mammalia | *Hyaena brunnea* | Brown hyena | 42250 | 85 | | | | | 12 | | | 235 |
| Mammalia | *Hylobates lar* | White-handed gibbon | 6810 | 64 | | | | | 20 | | 0.002 | 254 |
| Mammalia | *Hylobates syndactylus* | Siamang | 10900 | 85 | | | | | 25 | | | 129 |
| Mammalia | *Ictonyx striatus* | Striped polecat | 910 | 64 | | | | | 5 | | 0.0361 | 206 |
| Mammalia | *Isoodon macrourus* | Northern brown bandicoot | 1551 | 64 | | | | | 2 | 713.03 | 0.0145 | 310 |

| | | | | | | | | | | | | |
|---|---|---|---|---|---|---|---|---|---|---|---|---|
| Mammalia | *Isoodon obesulus* | Southern brown bandicoot | 717 | 38 | 700 | 64 | | | 3.5 | 339.6069 | 0.0135 | 215 |
| Mammalia | *Kobus ellipsiprymnus* | Waterbuck | 175333 | 85 | | | | | 18.5 | | | 297 |
| Mammalia | *Lagorchestes conspicillatus* | Spectacled hare-wallaby | 2660 | 85 | | | | | 6 | | | 227 |
| Mammalia | *Lemur catta* | Ring-tailed lemur | 2555 | 85 | | | | | 27.1 | | | 184 |
| Mammalia | *Leontopithecus rosalia* | Golden lion tamarin | 654.5 | 64 | | | | | 12 | | 0.0077 | 200 |
| Mammalia | *Lepus americanus* | Snowshoe hare | 1603 | 85 | | | | | 5 | | | 289 |
| Mammalia | *Lepus europaeus* | European hare | 4175 | 64 | | | | | 13 | | 0.0191 | 119 |
| Mammalia | *Liomys adspersus* | Spiny pocket mouse | 42 | 85 | | | | | 1.5 | | | 159 |
| Mammalia | *Litocranius walleri* | Gerenuk | 43500 | 85 | | | | | 8 | | | 210 |
| Mammalia | *Loxodonta africana* | African bush elephant | 5E+06 | 64 | | | | | 65 | | 0.0003 | 238 |
| Mammalia | *Lutreolina crassicaudata* | Lutrine opossum | 812 | 38 | | | | | 3 | 509.6935 | | 152 |
| Mammalia | *Lycaon pictus* | African hunting dog | 8750 | 85 | | | | | 11 | | | 161 |
| Mammalia | *Lynx lynx* | Eurasian lynx | 23000 | 64 | | | | | 17 | | 0.0097 | 148 |

| | | | | | | | | | | | | |
|---|---|---|---|---|---|---|---|---|---|---|---|---|
| Mammalia | *Macaca arctoides* | Stump-tailed macaque | 9275 | 85 | | | | | 30 | | | 184 |
| Mammalia | *Macaca mulatta* | Rhesus macaque | 8235 | 64 | | | | | 28 | | 0.0012 | 242 |
| Mammalia | *Macaca sinica* | Toque macaque | 4370 | 85 | | | | | 30 | | | 271 |
| Mammalia | *Macaca sylvanus* | Barbary ape | 10875 | 85 | | | | | 22 | | | 271 |
| Mammalia | *Macropus fuliginosus* | Western gray kangaroo | 28250 | 64 | | | | | 20 | | 0.0108 | 85 |
| Mammalia | *Macropus giganteus* | Eastern gray kangaroo | 49500 | 64 | | | | | 20 | | 0.0066 | 85 |
| Mammalia | *Macropus parma* | Parma wallaby | 4250 | 64 | | | | | 8 | | 0.01 | 85 |
| Mammalia | *Macropus robustus* | Wallaroo | 30000 | 64 | 29648 | 85 | | | 18.5 | | 0.0038 | 188 |
| Mammalia | *Macropus rufogriseus* | Red-necked wallaby | 16200 | 85 | | | | | 18.5 | | | 223 |
| Mammalia | *Macropus rufus* | Red kangaroo | 55000 | 85 | | | 28500 | 38 | 22 | | | 188 |
| Mammalia | *Macrotis lagotis* | Rabbit-bandicoot | 1246 | 85 | | | | | 7.2 | 613.19 | | 135 |
| Mammalia | *Madoqua kirkii* | Kirk's dik-dik | 5000 | 64 | | | | | 5 | | 0.0124 | 164 |
| Mammalia | *Marmosa murina* | Murine opossum | 26 | 85 | | | | | 1 | | | 192 |

| | | | | | | | | | | | | |
|---|---|---|---|---|---|---|---|---|---|---|---|---|
| Mammalia | *Marmosa robinsoni* | Robinson's mouse opossum | 122 | 38 | | | | | 2 | 147.5895 | | 135 |
| Mammalia | *Marmota marmota* | Apline marmot | 3500 | 85 | | | | | 15 | | | 245 |
| Mammalia | *Martes americana* | American marten | 966.5 | 85 | | | 900 | 38 | 13 | 597.9833 | | 255 |
| Mammalia | *Martes pennanti* | Fisher | 3175 | 85 | | | | | 10 | | | 261 |
| Mammalia | *Meles meles* | Old world badger | 13000 | 64 | | | | | 15 | | 0.0196 | 220 |
| Mammalia | *Mellivora capensis* | Honey badger | 10000 | 85 | | | | | 26.4 | | | 196 |
| Mammalia | *Mephitis mephitis* | Striped skunk | 3500 | 64 | | | | | 6 | | 0.0143 | 126 |
| Mammalia | *Meriones unguiculatus* | Mongolian jird | 53.2 | 64 | | | | | 3 | 75.47 | 0.0324 | 305 |
| Mammalia | *Metachirus nudicaudatus* | Brown four-eyed opossum | 336 | 38 | | | | | 2 | 279.4945 | | 135 |
| Mammalia | *Microcebus murinus* | Gray mouse lemur | 64.8 | 64 | | | | | 10 | | 0.0362 | 181 |
| Mammalia | *Microcebus rufus* | Brown mouse lemur | 55.5 | 85 | | | | | 8 | | | 321 |
| Mammalia | *Micromys minutus* | Harvest mouse | 7.6 | 85 | | | 7.37 | 38 | 1 | 21.078 | | 306 |
| Mammalia | *Microtus agrestis* | Field vole | 25 | 85 | | | 28 | 38 | 1.66 | 66.28537 | | 241 |
| Mammalia | *Microtus arvalis* | Common vole | 27.5 | 64 | 21.9 | 85 | | | 1.66 | 68.75 | 0.0561 | 241 |

| | | | | | | | | | | | | |
|---|---|---|---|---|---|---|---|---|---|---|---|---|
| Mammalia | *Microtus guentheri* | Vole | 43.8 | 38 | | | | | 0.5 | 77.70604 | | 134 |
| Mammalia | *Microtus ochrogaster* | Prairie vole | 48.2 | 85 | 40 | 64 | 46.7 | 38 | 1 | 79.62561 | 0.0666 | 176 |
| Mammalia | *Microtus pennsylvanicus* | Meadow vole | 38.2 | 85 | | | 38.9 | 38 | 1 | 71.29453 | | 284 |
| Mammalia | *Microtus subterraneus* | European pine vole | 17.8 | 85 | | | | | 1.75 | | | 144 |
| Mammalia | *Monodelphis brevicaudata* | Short bare-tailed opossum | 91.5 | 85 | | | 75.5 | 38 | 1 | 88.60531 | | 247 |
| Mammalia | *Muscardinus avellanarius* | Hazel dormouse | 23.5 | 85 | | | | | 4 | 79.065 | | 197 |
| Mammalia | *Mustela erminea* | Ermine | 125.5 | 85 | | | 75 | 38 | 4.5 | 139.8433 | | 153 |
| Mammalia | *Mustela lutreola* | European mink | 590 | 85 | | | | | 10 | | | 244 |
| Mammalia | *Mustela putorius* | European polecat | 809 | 64 | | | | | 6 | | 0.0328 | 85 |
| Mammalia | *Mustela vison* | American mink | 660 | 38 | 925 | 64 | | | 10 | 440.4273 | 0.0266 | 176 |
| Mammalia | *Myodes glareolus* | Bank vole | 21.5 | 85 | | | | | 2 | | | 176 |
| Mammalia | *Nannospalax spp.* | Mediterranean blind mole rat | 160 | 85 | | | | | 4.5 | 134.94 | | 275 |
| Mammalia | *Napaeozapus insignis* | Woodland jumping mouse | 22 | 38 | | | | | 4 | 43.91335 | | 245 |
| Mammalia | *Neotoma floridana* | Eastern woodrat | 291 | 64 | | | | | 3 | | 0.0654 | 100 |

| | | | | | | | | | | | | |
|---|---|---|---|---|---|---|---|---|---|---|---|---|
| Mammalia | *Nycticebus coucang* | Slow loris | 1129 | 85 | | | 1160 | 38 | 14 | 358.5209 | | 245 |
| Mammalia | *Ochotona collaris* | Collared pika | 129 | 85 | | | | | 6 | | | 176 |
| Mammalia | *Ochotona curzoniae* | Plateau pika | 109 | 85 | | | | | 4.25 | | | 265 |
| Mammalia | *Ochotona princeps* | North American pika | 109 | 85 | 100 | 64 | | | 4.3 | | 0.0376 | 266 |
| Mammalia | *Ochrotomys nuttalli* | Golden mouse | 19.5 | 38 | | | | | 2.5 | 32.04717 | | 258 |
| Mammalia | *Octodon degus* | Degu | 235 | 64 | 199.6 | 85 | | | 4 | 177.43 | 0.0323 | 298 |
| Mammalia | *Odocoileus hemionus* | Mule deer | 57000 | 64 | | | | | 22 | | 0.0057 | 85 |
| Mammalia | *Odocoileus virginianus* | White-tailed deer | 87000 | 64 | | | | | 19 | | 0.007 | 217 |
| Mammalia | *Ondatra zibethicus* | Muskrat | 1005 | 38 | | | | | 3 | 684.0912 | | 245 |
| Mammalia | *Ornithorhynchus anatinus* | Duck-billed platypus | 1030 | 85 | 1030 | 85 | 693 | 38 | 17 | 293.4248 | | 112 |
| Mammalia | *Orycteropus afer* | Aardvark | 48000 | 38 | | | | | 23 | 8822.792 | | 115 |
| Mammalia | *Oryctolagus cuniculus* | Old World rabbit | 2168 | 85 | | | | | 9 | | | 176 |
| Mammalia | *Otomys irroratus* | Southern African vlei rat | 102 | 38 | | | | | 2 | 88.44719 | | 102 |
| Mammalia | *Ovibos moschatus* | Muskox | 315000 | 64 | | | | | 23 | | 0.0041 | 122 |
| Mammalia | *Ovis canadensis* | Bighorn sheep | 70275 | 64 | 67031 | 85 | | | 19 | | 0.0031 | 110 |

| | | | | | | | | | | | | |
|---|---|---|---|---|---|---|---|---|---|---|---|---|
| Mammalia | *Pan troglodytes* | Chimpanzee | 44984 | 64 | | | | | 55 | | 0.0007 | 186 |
| Mammalia | *Panthera leo* | Lion | 98000 | 38 | 175000 | 64 | | | 15 | 17130.2 | 0.0035 | 219 |
| Mammalia | *Panthera pardus* | Leopard | 53750 | 64 | | | | | 23.2 | | 0.0079 | 104 |
| Mammalia | *Panthera tigris* | Tiger | 137900 | 38 | | | | | 26 | 25267.61 | | 224 |
| Mammalia | *Papio cynocephalus* | Yellow baboon | 23000 | 85 | | | | | 40 | | | 271 |
| Mammalia | *Papio hamadryas* | Hamadryas baboon | 18000 | 64 | | | | | 27 | | 0.0026 | 85 |
| Mammalia | *Perameles gunni* | Eastern barred bandicoot | 695 | 85 | 766 | 64 | 837 | 38 | 3 | 561.2457 | 0.0142 | 281 |
| Mammalia | *Perameles nasuta* | Long-nosed bandicoot | 645 | 38 | | | | | 3 | 384.6524 | | 135 |
| Mammalia | *Perodicticus potto* | Potto | 1225 | 64 | 968.6 | 85 | | | 10 | 399.55 | 0.0125 | 182 |
| Mammalia | *Perognathus longimembris* | Little pocket mouse | 8.5 | 85 | 8 | 64 | 8.9 | 38 | 4.3 | 13.39517 | 0.0766 | 162 |
| Mammalia | *Perognathus parvus* | Great basin pocket mouse | 19.2 | 85 | | | | | 4 | | | 248 |
| Mammalia | *Peromyscus leucopus* | White-footed mouse | 22.3 | 85 | 23 | 64 | 26 | 38 | 2.1 | 47.82757 | 0.0456 | 133 |
| Mammalia | *Peromyscus maniculatus* | Deer mouse | 20.5 | 64 | | | | | 2 | 42.85 | 0.0643 | 85 |
| Mammalia | *Peromyscus polionotus* | Oldfield mouse | 14 | 64 | 12 | 85 | | | 1 | | 0.0558 | 143 |

| | | | | | | | | | | | | |
|---|---|---|---|---|---|---|---|---|---|---|---|---|
| Mammalia | *Phascogale tapoatafa* | Brush-tailed phascogale | 153.7 | 85 | | | 157 | 38 | 3.5 | 135.3086 | | 277 |
| Mammalia | *Phascolarctos cinereus* | Koala | 4765 | 38 | 9300 | 64 | | | 13 | 1298.093 | 0.0028 | 219 |
| Mammalia | *Phenacomys intermedius* | Western heather vole | 21.5 | 38 | | | | | 4 | 67.99437 | | 226 |
| Mammalia | *Philander opossum* | Gray four-eyed opossum | 751 | 38 | | | | | 2.5 | 424.2633 | | 128 |
| Mammalia | *Planigale ingrami* | Long-tailed planigale | 8.8 | 85 | | | | | 1.3 | | | 135 |
| Mammalia | *Planigale tenuirostris* | Narrow-nosed planigale | 7.1 | 38 | | | | | 3 | 16.21102 | | 268 |
| Mammalia | *Poliocitellus franklini* | Franklin's ground squirrel | 607 | 85 | | | | | 5 | | | 252 |
| Mammalia | *Pongo pygmaeus* | Orangutan | 64475 | 64 | | | | | 58 | | 0.0009 | 313 |
| Mammalia | *Potamochoerus porcus* | Red river hog | 88000 | 64 | | | | | 15 | | 0.0099 | 209 |
| Mammalia | *Potorous tridactylus* | Long-nosed potoroo | 1430 | 64 | 1045.5 | 85 | 976 | 38 | 7 | 520.0876 | 0.0122 | 177 |
| Mammalia | *Priodontes maximus* | Giant armadillo | 45190 | 38 | | | | | 15 | 4771.811 | | 232 |
| Mammalia | *Procavia capensis* | Rock hyrax | 2400 | 38 | 3600 | 64 | | | 8.5 | 731.8892 | 0.0112 | 188 |
| Mammalia | *Procyon cancivorus* | Crab-eating raccoon | 1160 | 85 | | | | | 16 | | | 218 |

| | | | | | | | | | | | | |
|---|---|---|---|---|---|---|---|---|---|---|---|---|
| Mammalia | *Procyon lotor* | Common raccoon | 5075 | 38 | 6000 | 64 | | | 16 | 1598.625 | 0.0153 | 196 |
| Mammalia | *Proechimys semispinosus* | Spiny rat | 498 | 38 | 500 | 64 | | | 4.4 | 316.9601 | 0.0131 | 249 |
| Mammalia | *Psammomys obesus* | Fat sand rat | 212 | 64 | | | | | 3 | | 0.0304 | 116 |
| Mammalia | *Pseudocheirus peregrinus* | Ring-tailed possum | 916 | 38 | | | | | 5 | 458.0722 | | 190 |
| Mammalia | *Puma concolor* | Cougar | 63000 | 64 | 37200 | 85 | | | 9 | | 0.0061 | 165 |
| Mammalia | *Rangifer tarandus* | Caribou | 101250 | 64 | 85000 | 85 | | | 4.6 | | 0.0047 | 240 |
| Mammalia | *Rattus fuscipes* | Bush rat | 76 | 38 | | | | | 3.8 | 89.29607 | | 303 |
| Mammalia | *Rattus lutreolus* | Australian swamp rat | 109 | 38 | 115 | 64 | | | 2.4 | 72.31488 | 0.0277 | 303 |
| Mammalia | *Rattus norvegicus* | Norway rat | 206.9 | 85 | | | | | 3 | | | 132 |
| Mammalia | *Rattus rattus* | Rat | 200 | 64 | 117 | 85 | | | 3 | | 0.0207 | 132 |
| Mammalia | *Redunca fulvorufula* | Mountain reedbuck | 30000 | 64 | | | | | 12 | | 0.0093 | 164 |
| Mammalia | *Reithrodontomys megalotis* | Western harvest mouse | 9 | 38 | | | | | 1.5 | 25.47207 | | 158 |
| Mammalia | *Reithrodontomys spp.* | American harvest mouse | 10.3 | 85 | | | | | 1.5 | | | 158 |
| Mammalia | *Rhabdomys pumilio* | Four-striped grass mouse | 39.6 | 38 | 51 | 64 | | | 1.6 | 35.56981 | 0.0361 | 145 |
| Mammalia | *Rhombomys opimus* | Great gerbil | 285 | 85 | | | | | 4 | | | 242 |

| | | | | | | | | | | | | |
|---|---|---|---|---|---|---|---|---|---|---|---|---|
| Mammalia | *Saguinus nigricollis* | Black-mantled tamarin | 519.2 | 64 | | | | | 13 | | 0.0079 | 182 |
| Mammalia | *Saguinus oedipus* | Cotton-top tamarin | 445.5 | 64 | | | | | 13 | | 0.0102 | 182 |
| Mammalia | *Sarcophilus harrisii* | Tasmanian devil | 5775 | 38 | | | | | 7 | 1672.487 | | 219 |
| Mammalia | *Scalopus aquaticus* | Eastern aquatic mole | 48 | 38 | | | | | 3 | 83.22678 | | 183 |
| Mammalia | *Sciurus carolinensis* | Eastern gray squirrel | 440 | 38 | 533 | 64 | | | 12.5 | 343.7961 | 0.0234 | 196 |
| Mammalia | *Sciurus niger* | Eastern fox squirrel | 800 | 64 | | | | | 12.6 | | 0.018 | 203 |
| Mammalia | *Sciurus vulgaris* | Eurasian red squirrel | 600 | 64 | | | | | 12 | | 0.021 | 85 |
| Mammalia | *Setonix brachyurus* | Quokka | 2702 | 85 | | | | | 10 | | | 287 |
| Mammalia | *Sicista betulina* | Northern birch mouse | 10 | 85 | | | | | 3 | | | 263 |
| Mammalia | *Sicista sp.* | Birch mouse | 10 | 85 | | | | | 3.3 | | | 193 |
| Mammalia | *Sigmodon hispidus* | Hispid cotton mouse | 159.6 | 85 | 185 | 64 | 139.3 | 38 | 2.5 | 228.0326 | 0.0362 | 111 |
| Mammalia | *Sminthopsis crassicaudata* | Fat-tailed dunnart | 16.4 | 38 | | | | | 1.3 | 33.6602 | | 135 |
| Mammalia | *Sminthopsis psammophila* | Sandhill dunnart | 55 | 85 | | | | | 2 | | | 221 |
| Mammalia | *Sorex alpinus* | Alpine shrew | 7.8 | 85 | | | | | 1.4 | | | 176 |

| | | | | | | | | | | | | |
|---|---|---|---|---|---|---|---|---|---|---|---|---|
| Mammalia | *Sorex araneus* | Common shrew | 8.4 | 85 | | | | | 1.5 | | | 131 |
| Mammalia | *Sorex cinereus* | Cinereus shrew | 5.2 | 85 | | | | | 1.8 | | | 154 |
| Mammalia | *Sorex coronatus* | Crowned shrew | 9.1 | 85 | | | | | 1.7 | | | 301 |
| Mammalia | *Sorex minutus* | Eurasian pygmy shrew | 4.1 | 85 | | | | | 1.8 | | | 257 |
| Mammalia | *Sorex vagrans* | Vagrant shrew | 5.2 | 85 | | | | | 1.3 | | | 154 |
| Mammalia | *Spalax ehrenbergi* | Palestine mole rat | 133.8 | 85 | | | 135.3 | 38 | 3 | 163.5204 | | 156 |
| Mammalia | *Spermophilus citellus* | European ground squirrel | 240 | 38 | | | | | 6 | 240.0963 | | 187 |
| Mammalia | *Spermophilus lateralis* | Ground squirrel | 240 | 38 | | | | | 11 | 125.0856 | | 285 |
| Mammalia | *Spermophilus sp.* | Golden-mantled ground squirrel | 157.6 | 64 | 249.6 | 85 | | | 7 | 170.96 | 0.0411 | 106 |
| Mammalia | *Suncus etruscus* | Etruscan shrew | 2.4 | 85 | 2.1 | 64 | | | 1 | | 0.174 | 279 |
| Mammalia | *Suncus murinus* | Asian house shrew | 39.7 | 85 | 45.9 | 64 | | | 3 | | 0.0643 | 286 |
| Mammalia | *Sundamys muelleri* | Giant sunda rat | 334 | 85 | | | | | 0.5 | | | 231 |
| Mammalia | *Suricata suricata* | Meerkat | 850 | 38 | 776 | 64 | | | 15 | 369.8655 | 0.0161 | 309 |

| | | | | | | | | | | | | |
|---|---|---|---|---|---|---|---|---|---|---|---|---|
| Mammalia | *Sus scrofa* | Wild boar | 180000 | 64 | 135000 | 85 | | | 10 | | 0.0095 | 194 |
| Mammalia | *Sylvilagus audubonii* | Audubon's cottontail rabbit | 686.9 | 85 | | | | | 2 | | | 213 |
| Mammalia | *Sylvilagus floridanus* | Eastern cottontail | 1150 | 85 | 1530 | 64 | | | 5 | | 0.0243 | 280 |
| Mammalia | *Syncerus caffer* | African buffalo | 700000 | 64 | | | | | 18 | | 0.0026 | 85 |
| Mammalia | *Tachyglossus aculeatus* | Short-beaked echidna | 4250 | 64 | 2909 | 85 | | | 48 | 915.81 | 0.013 | 101 |
| Mammalia | *Tamias minimus* | Least chipmunk | 49.3 | 85 | | | 45.8 | 38 | 0.66 | 80.65219 | | 311 |
| Mammalia | *Tamias striatus* | Eastern American chipmunk | 87.4 | 38 | | | | | 3 | 88.63667 | | 236 |
| Mammalia | *Tamias townsendii* | Townsend's chipmunk | 75 | 64 | | | | | 3 | | 0.0333 | 236 |
| Mammalia | *Tamiasciurus hudsonicus* | Red squirrel | 219.6 | 85 | 200 | 64 | 228.3 | 38 | 7 | 238.0098 | 0.0216 | 225 |
| Mammalia | *Tarsipes rostratus* | Honey possum | 10 | 38 | | | | | 1.5 | 33.5166 | | 315 |
| Mammalia | *Tarsius tarsier* | Spectral tarsier | 173 | 85 | | | | | 10.75 | | | 283 |
| Mammalia | *Taurotragus oryx* | Eland | 500000 | 64 | 141404 | 85 | | | 16 | | 0.0019 | 125 |
| Mammalia | *Taxidea taxus* | American badger | 9000 | 38 | | | | | 14 | 2700 | | 196 |
| Mammalia | *Thomomys bottae* | Botta's pocket gopher | 143 | 38 | | | | | 4.5 | 147.7128 | | 142 |

| | | | | | | | | | | | | |
|---|---|---|---|---|---|---|---|---|---|---|---|---|
| Mammalia | *Thomomys talpoides* | Pocket gopher | 106.8 | 38 | | | | | 1.6 | 171.0982 | | 115 |
| Mammalia | *Thylogale billardierii* | Tasmanian pademelon | 5000 | 64 | | | | | 7 | | 0.0098 | 229 |
| Mammalia | *Tolypeutes matacus* | Three-banded armadillo | 1160 | 38 | | | | | 15 | 352.0817 | | 232 |
| Mammalia | *Tragelaphus imberbis* | Lesser kudu | 82500 | 85 | | | | | 10 | | | 210 |
| Mammalia | *Tragelaphus scriptus* | Bushbuck | 60000 | 64 | | | | | 14 | | 0.0044 | 164 |
| Mammalia | *Tragelaphus spekei* | Sitatunga | 87500 | 85 | | | | | 20 | | | 198 |
| Mammalia | *Trichosurus caninus* | Mountain brush-tailed possum | 3150 | 85 | | | | | 17 | | | 299 |
| Mammalia | *Trichosurus vulpecula* | Brush-tailed possum | 2005 | 38 | 2395 | 64 | | | 10 | 899.7082 | 0.0206 | 133 |
| Mammalia | *Tupaia glis* | Common tree shrew | 123 | 38 | | | | | 4 | 103.6621 | | 205 |
| Mammalia | *Urocitellus richardsonii* | Richardson's ground squirrel | 266.3 | 85 | | | | | 6 | | | 233 |
| Mammalia | *Ursus americanus* | American black bear | 154250 | 64 | | | | | 33 | | 0.0029 | 85 |
| Mammalia | *Ursus arctos* | Grizzly bear | 277500 | 85 | | | | | 25 | | | 139 |
| Mammalia | *Ursus maritimus* | Polar bear | 475000 | 64 | | | | | 30 | | 0.0022 | 146 |
| Mammalia | *Vicugna vicugna* | Vicugna | 50000 | 64 | | | | | 13 | | 0.0057 | 136 |
| Mammalia | *Vulpes lagopus* | Arctic fox | 3600 | 85 | | | | | 10 | | | 96 |
| Mammalia | *Vulpes velox* | Swift fox | 1769 | 85 | 2400 | 64 | | | 9 | | 0.0315 | 270 |

| | | | | | | | | | | | | |
|---|---|---|---|---|---|---|---|---|---|---|---|---|
| Mammalia | *Vulpes vulpes* | Red fox | 4440 | 38 | 8500 | 64 | | | 15 | 2271.51 | 0.0177 | 239 |
| Mammalia | *Xerospermophilus mohavensis* | Mohave ground squirrel | 240 | 85 | | | | | 5 | 125.0865 | | 178 |
| Mammalia | *Zapus hudsonicus* | Meadow jumping mouse | 26.1 | 85 | 18 | 64 | 23.8 | 38 | 3 | 38.37949 | 0.0328 | 191 |
| Mammalia | *Zapus princeps* | Western jumping mouse | 21 | 85 | | | | | 4 | | | 121 |
| Reptilia | *Acanthodactylus boskianus* | | 7.8 | 38 | 19.6 | 62 | | | 9 | 5.572925 | | 62 |
| Reptilia | *Acanthodactylus dumerilii* | | 7.3 | 62 | | | | | 1.5 | | | 62 |
| Reptilia | *Acanthodactylus erythrurus* | | 9 | 38 | 13.7 | 62 | | | 2.1 | 4.881291 | | 62 |
| Reptilia | *Acanthodactylus pardalis* | | 9.7 | 38 | 10.6 | 62 | | | 4 | 7.383813 | | 62 |
| Reptilia | *Acontias meleagris* | | 7.3 | 38 | | | | | 3.9 | 1.28466 | | 160 |
| Reptilia | *Alligator mississippiensis* | | 150000 | 85 | 1287 | 38 | | | 73.1 | 120.01 | 0.00017 | 314 |
| Reptilia | *Alligator sinensis* | Chinese alligator | 35800 | 93 | | | | | 50 | | | 93 |
| Reptilia | *Alsophylax pipiens* | | 1.7 | 62 | | | | | 3.5 | | | 62 |
| Reptilia | *Amalosia lesueurii* | | 10.3 | 62 | | | | | 9.5 | | | 62 |
| Reptilia | *Amblyrhynchus cristatus* | | 7406.5 | 62 | | | | | 28 | 64.81586 | | 62 |

| | | | | | | | | | | | | |
|---|---|---|---|---|---|---|---|---|---|---|---|---|
| Reptilia | *Amphibolurus muricatus* | | 46.8 | 62 | | | | | 9.9 | 6.105519 | | 117 |
| Reptilia | *Amphibolurus norrisi* | | 38.2 | 62 | | | | | 7 | | | 62 |
| Reptilia | *Anatololacerta anatolica* | | 9.8 | 62 | | | | | 10 | | | 62 |
| Reptilia | *Anolis acutus* | | 5 | 62 | 1.481 | 65 | | | 1.6 | 2.58 | 0.002 | 62 |
| Reptilia | *Anolis carolinensis* | | 10.5 | 62 | | | | | 7.1 | 2.061847 | | 117 |
| Reptilia | *Anolis cristatellus* | | 9.8 | 62 | | | | | 6.9 | | | 62 |
| Reptilia | *Anolis cupreus* | | 3.9 | 62 | | | | | 1.5 | | | 62 |
| Reptilia | *Anolis garmani* | | 52.6 | 62 | | | | | 10 | | | 62 |
| Reptilia | *Anolis gemmosus* | | 6 | 62 | | | | | 3 | | | 62 |
| Reptilia | *Anolis gundlachi* | | 9.1 | 62 | | | | | 3 | | | 62 |
| Reptilia | *Anolis humilis* | | 2.7 | 62 | | | | | 1.5 | | | 62 |
| Reptilia | *Anolis intermedius* | | 3.3 | 62 | | | | | 2 | | | 62 |
| Reptilia | *Anolis limifrons* | | 1.4 | 38 | | | | | 1 | 0.3981 | | 62 |
| Reptilia | *Anolis stratulus* | | 4.8 | 62 | | | | | 1.6 | | | 62 |
| Reptilia | *Anolis tropidolepis* | | 4.3 | 62 | | | | | 2 | | | 62 |
| Reptilia | *Aspidoscelis deppei* | | 23.7 | 62 | | | | | 2 | | | 62 |
| Reptilia | *Aspidoscelis dixoni* | | 40 | 62 | | | | | 4 | | | 62 |
| Reptilia | *Aspidoscelis exsanguis* | | 30.7 | 62 | | | | | 4 | | | 62 |

| | | | | | | | | | | | | |
|---|---|---|---|---|---|---|---|---|---|---|---|---|
| Reptilia | *Aspidoscelis laredoensis* | | 21.4 | 62 | | | | | 3 | | | 62 |
| Reptilia | *Aspidoscelis neomexicana* | | 18.6 | 62 | | | | | 4 | | | 62 |
| Reptilia | *Aspidoscelis rodecki* | | 12.1 | 62 | | | | | 2 | | | 62 |
| Reptilia | *Aspidoscelis scalaris* | | 59.5 | 62 | | | | | 3 | | | 62 |
| Reptilia | *Aspidoscelis tesselata* | | 36.7 | 62 | | | | | 5 | | | 62 |
| Reptilia | *Aspidoscelis tigris* | | 79.1 | 62 | | | | | 8 | | | 62 |
| Reptilia | *Aspidoscelis uniparens* | | 18.6 | 62 | | | | | 4 | | | 62 |
| Reptilia | *Basiliscus basiliscus* | | 554 | 62 | | | | | 7 | | | 62 |
| Reptilia | *Bassiana duperreyi* | | 10.5 | 62 | | | | | 7 | | | 62 |
| Reptilia | *Blanus cinereus* | | 2.4 | 38 | 16 | 62 | | | 0.7 | 1.09 | | 160 |
| Reptilia | *Boa constrictor* | Common boa constrictor | 35283 | 62 | 7815.5 | 38 | | | 40.4 | 273.6353 | 0.00003 | 62 |
| Reptilia | *Boaedon fuliginosus* | | 315.1 | 62 | | | | | 9 | | | 62 |
| Reptilia | *Caimen latirostris* | Broad-nosed caiman | 60000 | 62 | | | | | 22 | | | 294 |
| Reptilia | *Callisaurus draconoides* | | 39.7 | 62 | | | | | 5.9 | | 0.00518 | 62 |
| Reptilia | *Caretta caretta* | | 57778 | 85 | | | | | | | 1.33E-05 | |

| | | | | | | | | | | | | |
|---|---|---|---|---|---|---|---|---|---|---|---|---|
| Reptilia | *Carlia bicarinata* | | 2 | 62 | | | | | 2 | | | 62 |
| Reptilia | *Carlia rostralis* | | 6.8 | 62 | | | | | 3.8 | | | 62 |
| Reptilia | *Carlia rubrigularis* | | 4.4 | 62 | | | | | 3.6 | | | 62 |
| Reptilia | *Carlia storri* | | 1.8 | 62 | | | | | 3.3 | | | 62 |
| Reptilia | *Chalcides chalcides* | | 34.2 | 62 | | | | | 4 | | | 62 |
| Reptilia | *Chalcides ocellatus* | | 22.06 | 38 | 218.6 | 62 | | | 8.9 | 5.11 | | 160 |
| Reptilia | *Chelonia mydas* | Green sea turtle | 160000 | 85 | 50889 | 65 | | | 62 | | 3.41E-05 | 171 |
| Reptilia | *Chelydra serpentina* | Common snapping turtle | 10250 | 85 | 3473 | 38 | | | 47 | 192.7 | | 62 |
| Reptilia | *Chioninia coctei* | | 1229.7 | 62 | | | | | 16 | | | 62 |
| Reptilia | *Chlamydosarus kingi* | Filled dragon | 620.8 | 62 | | | | | 6 | | | 173 |
| Reptilia | *Chrysemys picta* | Painted turtle | 350 | 85 | | | | | 61 | | 0.00034 | 136 |
| Reptilia | *Clemmys guttata* | Spotted turtle | 50.37 | 85 | | | | | 110 | | 0.00235 | 214 |
| Reptilia | *Cnemidophorus murinus* | | 85 | 38 | 125.7 | 62 | | | 5 | 13.07877 | | 62 |
| Reptilia | *Cnemidophorus sexlineatus* | Six-lined racerunner | 6 | 86 | | | | | 7.8 | | 0.01667 | 85 |
| Reptilia | *Cnemidophorus tigris* | | 26 | 86 | | | | | 7.8 | 5.34 | 0.00269 | 319 |
| Reptilia | *Coelognathus helena* | | 437.6 | 62 | | | | | 15 | | | 62 |
| Reptilia | *Coleonyx brevis* | | 5.4 | 62 | | | | | 5 | | | 62 |

| | | | | | | | | | | | | |
|---|---|---|---|---|---|---|---|---|---|---|---|---|
| Reptilia | *Coluber constrictor* | | 262 | 38 | 606.9 | 62 | | | 10 | 21.91 | 0.0006 | 62 |
| Reptilia | *Conolophus pallidus* | | 5288.6 | 62 | | | | | 60 | | | 62 |
| Reptilia | *Conolophus subcristatus* | | 6288.5 | 62 | | | | | 60 | | | 62 |
| Reptilia | *Cophosaurus texanus* | | 23.1 | 62 | | | | | 6 | | | 62 |
| Reptilia | *Coronella girondica* | | 66.5 | 62 | | | | | 16 | | | 62 |
| Reptilia | *Crocodylus moreletii* | Morelet's crocodiles | 51000 | 95 | | | | | 60 | | | 95 |
| Reptilia | *Crocodylus siamensis* | Siamese crocodile | 151000 | 86 | | | | | 80 | | | 89 |
| Reptilia | *Crotalus durissus* | Neotropical rattlesnake | 2883 | 62 | | | | | 19.8 | | | 62 |
| Reptilia | *Crotaphytus collaris* | | 30 | 38 | 68.1 | 62 | | | 10 | 10.6507 | | 62 |
| Reptilia | *Ctenophorus fordi* | | 4.4 | 62 | | | | | 2 | | | 62 |
| Reptilia | *Ctenophorus isolepis* | | 13.3 | 62 | | | | | 2 | | | 62 |
| Reptilia | *Ctenophorus maculosus* | | 7.9 | 62 | | | | | 3.5 | | | 62 |
| Reptilia | *Ctenophorus ornatus* | | 20.1 | 62 | | | | | 11 | | | 62 |
| Reptilia | *Ctenosaura hemilopha* | | 2724.7 | 62 | | | | | 13 | | | 62 |
| Reptilia | *Ctenotus atlas* | | 8.5 | 62 | | | | | 4 | | | 62 |
| Reptilia | *Ctenotus labillarderi* | Lancelin Island skink | 8.9 | 62 | | | | | 7 | 0.99 | | 294 |

| | | | | | | | | | | | | |
|---|---|---|---|---|---|---|---|---|---|---|---|---|
| Reptilia | *Ctenotus lancelini* | | 13.7 | 62 | | | | | 6 | | | 62 |
| Reptilia | *Ctenotus leonhardii* | | 2.8 | 38 | 10.1 | 62 | | | 7 | | | 62 |
| Reptilia | *Ctenotus regius* | | 11.8 | 62 | | | | | 4 | | | 62 |
| Reptilia | *Ctenotus schomburgkii* | | 11.8 | 62 | | | | | 5 | | | 62 |
| Reptilia | *Ctenotus strauchii* | | 4.8 | 62 | | | | | 2.5 | | | 62 |
| Reptilia | *Ctenotus taeniolatus* | | 14.8 | 62 | | | | | 5 | | | 62 |
| Reptilia | *Cyclura carinata* | | 5609.2 | 62 | | | | | 14 | | | 62 |
| Reptilia | *Cyclura cychlura* | | 11800 | 62 | | | | | 40 | | | 62 |
| Reptilia | *Cyclura nubila* | Cuban rock iguana | 17648 | 62 | | | | | 40 | | | 113 |
| Reptilia | *Darevskia armeniaca* | | 9 | 62 | | | | | 8 | | | 62 |
| Reptilia | *Darevskia brauneri* | | 8.3 | 62 | | | | | 13 | | | 62 |
| Reptilia | *Darevskia dahli* | | 6.1 | 62 | | | | | 6 | | | 62 |
| Reptilia | *Darevskia portschinskii* | | 7 | 62 | | | | | 6 | | | 62 |
| Reptilia | *Darevskia raddei* | | 10.2 | 62 | | | | | 6 | | | 62 |
| Reptilia | *Darevskia rostombekovi* | | 5.1 | 62 | | | | | 6 | | | 62 |
| Reptilia | *Darevskia rudis* | | 15.9 | 62 | | | | | 8 | | | 62 |
| Reptilia | *Darevskia unisexualis* | | 9 | 62 | | | | | 7 | | | 62 |
| Reptilia | *Darevskia valentini* | | 11.8 | 62 | | | | | 7 | | | 62 |

| | | | | | | | | | | | | |
|---|---|---|---|---|---|---|---|---|---|---|---|---|
| Reptilia | *Diadophis punctatus* | | 14.8 | 85 | | | | | 10 | 1.19 | 0.0002 | 114 |
| Reptilia | *Dinarolacerta mosorensis* | | 11.8 | 62 | | | | | 9 | | | 62 |
| Reptilia | *Diplodactylus conspicillatus* | | 5.5 | 62 | | | | | 3.2 | | | 62 |
| Reptilia | *Draco volans* | | 20.8 | 62 | | | | | 3 | | | 62 |
| Reptilia | *Egernia cunninghami* | | 414.9 | 62 | | | | | 20 | 45.22809 | | 169 |
| Reptilia | *Egernia kingii* | | 414.9 | 62 | | | | | 13 | | | 62 |
| Reptilia | *Elgaria coerulea* | | 45.7 | 62 | | | | | 8 | | | 62 |
| Reptilia | *Elgaria multicarinata* | | 116.5 | 62 | | | | | 15 | | | 62 |
| Reptilia | *Emoia atrocostata* | | 21.5 | 62 | | | | | 3.5 | | 0.00294 | 62 |
| Reptilia | *Eremias argus* | | 7.2 | 62 | | | | | 11 | | | 62 |
| Reptilia | *Eremias strauchi* | | 11.8 | 62 | | | | | 7 | | | 62 |
| Reptilia | *Ergenia whitii* | White's skink | 25 | 91 | | | | | 8.5 | | | 89 |
| Reptilia | *Eulamprus leuraensis* | | 12.7 | 62 | | | | | 6 | | | 62 |
| Reptilia | *Eulamprus tympanum* | | 21.6 | 62 | | | | | 15 | | | 62 |
| Reptilia | *Eumeces fasciatus* | | 13.2 | 62 | 22 | 66 | | | 8 | | 0.00136 | 138 |
| Reptilia | *Eutropis multifasciata* | | 63.8 | 62 | | | | | 5 | | | 62 |
| Reptilia | *Farancia abacura* | | 689.9 | 62 | | | | | 19 | | | 62 |
| Reptilia | *Furcifer labordi* | | 64.8 | 62 | | | | | 0.5 | | | 62 |

| | | | | | | | | | | | | |
|---|---|---|---|---|---|---|---|---|---|---|---|---|
| Reptilia | *Gallotia bravoana* | | 164 | 62 | | | | | 6 | | | 62 |
| Reptilia | *Gallotia intermedia* | | 91.5 | 62 | | | | | 10.4 | | | 62 |
| Reptilia | *Gallotia simonyi* | | 2671.6 | 62 | | | | | 20 | | | 62 |
| Reptilia | *Gambelia sila* | | 58.5 | 62 | | | | | 4.8 | | | 62 |
| Reptilia | *Gambelia wislizenii* | | 96.8 | 62 | | | | | 9.8 | | | 62 |
| Reptilia | *Gehyra variegata* | | 7.4 | 62 | | | | | 26 | | | 62 |
| Reptilia | *Gerrhonotus multicarinatus* | | 29 | 91 | | | | | 5 | 9.07957 | | 92 |
| Reptilia | *Gopherus agassizi* | | 20000 | 87 | 829.63 | 65 | | | 56 | | 0.0006 | 140 |
| Reptilia | *Heloderma suspectum* | Gila monster | 958.5 | 62 | | | | | | | | 62 |
| Reptilia | *Hemidactylus brookii* | | 12.2 | 62 | | | | | 4 | | | 62 |
| Reptilia | *Hemidactylus frenatus* | | 6.3 | 62 | | | | | 4 | 1.283539 | | 62 |
| Reptilia | *Hesperoedura reticulata* | | 7.2 | 62 | | | | | 11 | | | 62 |
| Reptilia | *Heterodon platirhinos* | | 183 | 62 | | | | | 9.1 | | 0.0025 | 62 |
| Reptilia | *Holbrookia maculata* | | 14.6 | 62 | | | | | 5 | | | 62 |
| Reptilia | *Holcosus quadrilineatus* | | 20 | 62 | | | | | 2 | | | 62 |
| Reptilia | *Homonota darwinii* | | 4.4 | 62 | | | | | 17 | | | 62 |

| | | | | | | | | | | | | |
|---|---|---|---|---|---|---|---|---|---|---|---|---|
| Reptilia | *Hoplodactylus duvaucelii* | | 87.6 | 62 | | | | | 50 | | | 62 |
| Reptilia | *Hoplodactylus maculatus* | Common gecko | 8 | 87 | | | | | 36 | | | 87 |
| Reptilia | *Iberolacerta aurelioi* | | 6.5 | 62 | | | | | 17 | | | 62 |
| Reptilia | *Iberolacerta monticola* | | 14 | 62 | | | | | 15 | | | 62 |
| Reptilia | *Iguana delicatissima* | | 3472.3 | 62 | | | | | 20 | | | 62 |
| Reptilia | *Iguana iguana* | | 1500 | 38 | 8220.7 | 62 | | | 41.5 | | 0.00014 | 98 |
| Reptilia | *Klauberina riversiana* | | 19 | 38 | | | | | 9.9 | 2.2 | | 157 |
| Reptilia | *Lacerta agilis* | | 8.4 | 38 | 33.6 | 62 | | | 12 | 6.295178 | | 62 |
| Reptilia | *Lacerta schreiberi* | | 55.4 | 62 | | | | | 10 | | | 62 |
| Reptilia | *Lacerta strigata* | | 91.5 | 62 | | | | | 4 | | | 62 |
| Reptilia | *Lacerta viridis* | European green lizard | 91.5 | 62 | | | | | 10 | 4.323572 | | 62 |
| Reptilia | *Lacerta vivipara* | vVviparous lizard | 12 | 86 | | | | | 12 | 2.472949 | | 103 |
| Reptilia | *Lamprolepis smaragdina* | | 34.8 | 62 | | | | | 5 | | | 62 |
| Reptilia | *Lampropholis delicata* | | 3.1 | 62 | | | | | 2 | | | 62 |
| Reptilia | *Lampropholis guichenoti* | | 2.6 | 62 | | | | | 5 | | | 62 |
| Reptilia | *Leposoma guianense* | | 1.2 | 62 | | | | | 2 | | | 62 |

| | | | | | | | | | | | | |
|---|---|---|---|---|---|---|---|---|---|---|---|---|
| Reptilia | *Liolaemus multiformis* | | 22 | 62 | | | | | 9 | | 0.00045 | 246 |
| Reptilia | *Liolaemus pictus* | | 37 | 62 | | | | | 9 | | | 62 |
| Reptilia | *Liolaemus quilmes* | | 22.9 | 62 | | | | | 7 | | | 62 |
| Reptilia | *Liopholis inornata* | | 12.7 | 62 | | | | | 10 | | | 62 |
| Reptilia | *Liopholis modesta* | | 31 | 62 | | | | | 5 | | | 62 |
| Reptilia | *Liopholis multiscutata* | | 18.9 | 62 | | | | | 10 | | | 62 |
| Reptilia | *Lissolepis coventryi* | | 79.7 | 62 | | | | | 9 | | | 62 |
| Reptilia | *Lucasium stenodactylum* | | 4.9 | 62 | | | | | 5 | | | 62 |
| Reptilia | *Lygodactylus capensis* | | 1.9 | 62 | | | | | 2.1 | | | 62 |
| Reptilia | *Macrochelys temminckii* | Alligator snapping turtle | 11300 | 85 | | | | | 45 | | | 94 |
| Reptilia | *Macroprotodon cucullatus* | | 25.7 | 62 | | | | | 16 | | | 62 |
| Reptilia | *Malpolon monspessulanus* | | 388.4 | 62 | | | | | 20 | | | 62 |
| Reptilia | *Mesalina olivieri* | | 3.3 | 62 | | | | | 5 | | | 62 |
| Reptilia | *Natrix erythrogaster* | Plain-bellied snake | 160 | 86 | | | | | 8.9 | | 0.00538 | 262 |
| Reptilia | *Natrix natrix* | | 37.5 | 38 | 2800.9 | 62 | | | 9 | 10.01764 | | 127 |

| | | | | | | | | | | | | |
|---|---|---|---|---|---|---|---|---|---|---|---|---|
| Reptilia | *Natrix tessellata* | | 283.8 | 62 | | | | | 14 | | | 62 |
| Reptilia | *Nephrurus stellatus* | | 16.6 | 62 | | | | | 8 | | | 62 |
| Reptilia | *Niveoscincus ocellatus* | | 12.7 | 62 | | | | | 12 | | | 62 |
| Reptilia | *Notechis scutatus* | | 1338.4 | 62 | | | | | 17 | | | 62 |
| Reptilia | *Oligosoma alani* | | 66.8 | 62 | | | | | 20 | | | 62 |
| Reptilia | *Oligosoma chloronoton* | | 44.3 | 62 | | | | | 8 | | | 62 |
| Reptilia | *Oligosoma lineoocellatum* | | 30.2 | 62 | | | | | 8 | | | 62 |
| Reptilia | *Oligosoma suteri* | | 45.4 | 62 | | | | | 12 | | | 62 |
| Reptilia | *Oligosoma whitakeri* | | 22.2 | 62 | | | | | 20 | | | 62 |
| Reptilia | *Ophisaurus apodus* | Armored glass lizard | 450 | 62 | | | | | 54 | | | 62 |
| Reptilia | *Ophisops elegans* | | 8 | 62 | | | | | 5 | | | 62 |
| Reptilia | *Osteolaemus tetraspis* | West African dwarf crocodile | 40000 | 62 | | | | | 34 | | | 151 |
| Reptilia | *Papuascincus stanleyanus* | | 4.1 | 62 | | | | | 6 | | | 62 |
| Reptilia | *Paralaudakia caucasia* | | 94.2 | 62 | | | | | 13 | | | 62 |
| Reptilia | *Paralaudakia stoliczkana* | | 143.4 | 62 | | | | | 10 | | | 62 |

| | | | | | | | | | | | | |
|---|---|---|---|---|---|---|---|---|---|---|---|---|
| Reptilia | *Parvilacerta parva* | | 5.6 | 62 | | | | | 8 | | | 62 |
| Reptilia | *Petrosaurus thalassinus* | | 141.1 | 62 | | | | | 20 | | | 62 |
| Reptilia | *Phrynocephalus guttatus* | | 6.3 | 62 | | | | | 5 | | | 62 |
| Reptilia | *Phrynocephalus mystaceus* | | 44.2 | 62 | | | | | 3.8 | | | 62 |
| Reptilia | *Phrynocephalus persicus* | | 4.7 | 62 | | | | | 5 | | | 62 |
| Reptilia | *Phrynoscma solace* | Regal horned lizard | 48 | 62 | | | | | 8 | | 0.005 | 230 |
| Reptilia | *Phrynosoma m'calli* | | 16 | 38 | 40 | 62 | | | 9 | 5.298626 | | 62 |
| Reptilia | *Phrynosoma platyrhinos* | | 27.5 | 62 | | | | | 8 | | | 62 |
| Reptilia | *Phymaturus tenebrosus* | | 55.9 | 62 | | | | | 16 | | | 62 |
| Reptilia | *Physignathus lesuerii* | | 504 | 38 | | | | | 6 | 80.6 | | 169 |
| Reptilia | *Plestiodon obsoletus* | Great Plains skink | 30 | 38 | 68.3 | 62 | | | 7.3 | | | 180 |
| Reptilia | *Plestiodon reynoldsi* | | 5.4 | 62 | | | | | 10 | | | 62 |
| Reptilia | *Plestiodon septentrionalis* | | 15.3 | 62 | | | | | 7 | | | 62 |
| Reptilia | *Plestiodon skiltonianus* | | 13.2 | 62 | | | | | 8 | | | 62 |
| Reptilia | *Podarcis bocagei* | | 8 | 62 | | | | | 4 | | | 62 |

| | | | | | | | | | | | | |
|---|---|---|---|---|---|---|---|---|---|---|---|---|
| Reptilia | *Podarcis erhardii* | | 13.7 | 62 | | | | | 5 | 1.533823 | | 62 |
| Reptilia | *Podarcis gaigeae* | | 22.9 | 62 | | | | | 5 | 3.600951 | | 62 |
| Reptilia | *Podarcis muralis* | | 5.5 | 38 | 11 | 62 | | | 5 | 4.236083 | | 105 |
| Reptilia | *Pogona vitticeps* | | 393.4 | 62 | | | | | 12 | | | 62 |
| Reptilia | *Pristidactylus achalensis* | | 33.9 | 62 | | | | | 11 | | | 62 |
| Reptilia | *Psammodromus hispanicus* | | 4.1 | 62 | | | | | 0.9 | | | 62 |
| Reptilia | *Psammophilus dorsalis* | | 66.2 | 62 | | | | | 4 | | | 62 |
| Reptilia | *Pseudemoia entrecasteauxii* | | 5.4 | 62 | | | | | 5 | | | 62 |
| Reptilia | *Ptyas mucosa* | | 3248.4 | 62 | | | | | 11.3 | | | 62 |
| Reptilia | *Ptyodactylus hasselquistii* | | 8.5 | 38 | 29.5 | 62 | | | 3.4 | 2.534776 | | 160 |
| Reptilia | *Python molurus* | | 33955 | 38 | 119935 | 62 | | | 34.2 | 807.38 | 0.0004475 | 62 |
| Reptilia | *Python regius* | | 830.5 | 38 | 1610 | 62 | | | 8.7 | 38.58 | | 160 |
| Reptilia | *Python reticulus* | | 14326 | 38 | | | | | 21.3 | 672.56 | 0.00045 | 160 |
| Reptilia | *Python sebae* | | 16140 | 38 | 50518 | 62 | | | 18 | 569.014 | 0.00096 | 160 |
| Reptilia | *Rhinechis scalaris* | | 386.6 | 62 | | | | | 16 | | | 62 |
| Reptilia | *Rhynchoedura ornata* | | 3.7 | 62 | | | | | 2.8 | | | 62 |
| Reptilia | *Sauromalus ater* | | 512.6 | 62 | | | | | 39 | | | 62 |
| Reptilia | *Sauromalus obesus* | | 150 | 62 | | | | | 39 | 21.83 | 0.000875 | 62 |
| Reptilia | *Scelarcis perspicillata* | | 5.1 | 62 | | | | | 5 | | | 62 |

| | | | | | | | | | | | | |
|---|---|---|---|---|---|---|---|---|---|---|---|---|
| Reptilia | *Sceloporus arenicolus* | | 12.1 | 62 | | | | | 2 | | | 62 |
| Reptilia | *Sceloporus bicanthalis* | | 8.8 | 62 | | | | | 1.3 | | | 62 |
| Reptilia | *Sceloporus cowlesi* | | 17.4 | 62 | | | | | 5 | | | 62 |
| Reptilia | *Sceloporus graciosus* | | 5 | 38 | 23.1 | 62 | | | 8 | 1.649415 | 0.00333 | 62 |
| Reptilia | *Sceloporus grammicus* | | 17.9 | 62 | | | | | 3 | | | 62 |
| Reptilia | *Sceloporus jarrovii* | | 36.9 | 62 | | | | | 2.8 | | | 62 |
| Reptilia | *Sceloporus magister* | | 80.7 | 62 | | | | | 6 | | 0.0021429 | 62 |
| Reptilia | *Sceloporus merriami* | | 10.4 | 62 | | | | | 6 | | | 62 |
| Reptilia | *Sceloporus occidentalis* | | 29.1 | 62 | | | | | 5 | 2.104958 | | 62 |
| Reptilia | *Sceloporus olivaceus* | | 13 | 38 | 56.1 | 62 | | | 3.8 | 8.902697 | | 62 |
| Reptilia | *Sceloporus scalaris* | | 16.2 | 62 | | | | | 5 | | | 62 |
| Reptilia | *Sceloporus tristichus* | | 23.8 | 62 | | | | | 5 | | | 62 |
| Reptilia | *Sceloporus undulatus* | | 3.8 | 38 | 24.5 | 62 | | | 4 | 1.4 | 0.003333 | 62 |
| Reptilia | *Sceloporus variabilis* | | 15.7 | 62 | | | | | 1.5 | 26.56 | | 62 |
| Reptilia | *Scincella lateralis* | | 3.9 | 62 | | | | | 4 | 0.6145 | | 62 |

| | | | | | | | | | | | | |
|---|---|---|---|---|---|---|---|---|---|---|---|---|
| Reptilia | *Sitana ponticeriana* | | 12.3 | 62 | | | | | 6 | | | 62 |
| Reptilia | *Sphaerodactylus vincenti* | | 1.6 | 62 | | | | | 4 | | | 62 |
| Reptilia | *Sphenodon punctatus* | | 430 | 38 | 1300 | 62 | | | 91 | 42.76 | 0.0001 | 62 |
| Reptilia | *Teira dugesii* | | 12.3 | 62 | | | | | 16 | | | 62 |
| Reptilia | *Telescopus fallax* | | 186.2 | 62 | | | | | 10 | | | 62 |
| Reptilia | *Teratoscincus roborowskii* | | 20.4 | 62 | | | | | 6 | | | 62 |
| Reptilia | *Thamnophis marcianus* | | 365.9 | 62 | | | | | 7 | | | 62 |
| Reptilia | *Thamnophis proximus* | | 31 | 38 | 562.3 | 62 | | | 9 | 10.48 | | 128 |
| Reptilia | *Thamnophis sirtalis* | Common garter snake | 150 | 38 | 862.3 | 62 | | | 14 | | 0.0005 | 308 |
| Reptilia | *Tiliqua rugosa* | Blue-tongued lizard | 617 | 38 | 1229.7 | 62 | | | 20.8 | 95.69247 | | 62 |
| Reptilia | *Tiliqua scincoides* | Australian blue-tongued lizard | 493 | 62 | | | | | 30 | 77.43 | | 202 |
| Reptilia | *Toropuku stephensi* | | 12.4 | 62 | | | | | 16 | | | 62 |
| Reptilia | *Trachemys scripta* | Common slider | 240 | 38 | | | | | 30 | | | 167 |
| Reptilia | *Trachylepis varia* | | 35.7 | 62 | | | | | 1.9 | | | 62 |
| Reptilia | *Trioceros hoehnelii* | | 29.8 | 62 | | | | | 4.5 | | | 62 |

| | | | | | | | | | | | | |
|---|---|---|---|---|---|---|---|---|---|---|---|---|
| Reptilia | *Tropidophis caymanensis* | | 46.9 | 62 | | | | | 25.1 | | | 62 |
| Reptilia | *Tropidurus hispidus* | | 87.7 | 62 | | | | | 2 | | | 62 |
| Reptilia | *Tropidurus torquatus* | | 79.2 | 62 | | | | | 3 | | | 62 |
| Reptilia | *Tupinambis nigropunctatus* | | 836 | 90 | | | | | | | | 90 |
| Reptilia | *Tympanocryptis centralis* | | 3.8 | 62 | | | | | 6 | | | 62 |
| Reptilia | *Uma inornata* | | 56.1 | 62 | | | | | 5 | | | 62 |
| Reptilia | *Uromastyx aegyptia* | | 1909.3 | 62 | 291.55 | 38 | | | 33 | 23.305 | | 62 |
| Reptilia | *Urosaurus graciosus* | | 11.2 | 62 | | | | | 5 | 23.305 | | 62 |
| Reptilia | *Urosaurus ornatus* | | 11.7 | 62 | | | | | 3 | | | 62 |
| Reptilia | *Uta stansburiana* | | 3.625 | 38 | 15.7 | 62 | | | 4.8 | 8.06 | 0.008 | 62 |
| Reptilia | *Varanus acanthurus* | | 73.61 | 86 | | | | | | | | |
| Reptilia | *Varanus exanthematicus* | | 1040 | 62 | | | | | 12.7 | | | 62 |
| Reptilia | *Varanus gouldi* | | 674 | 38 | | | | | 7.8 | 58.447 | | 160 |
| Reptilia | *Varanus griseus* | Desert monitor | 647 | 86 | 5542 | 62 | | | 14 | | 0.00069 | 290 |
| Reptilia | *Varanus komodoensis* | | 102478 | 62 | | | | | 62 | | | 62 |
| Reptilia | *Varanus niloticus* | Nile monitor | 1926 | 85 | | | | | 8.5 | | 0.0004798 | 121 |

| | | | | | | | | | | | | |
|---|---|---|---|---|---|---|---|---|---|---|---|---|
| Reptilia | *Varanus panoptes* | | 931 | 86 | | | | | | | | 277 |
| Reptilia | *Vipera berus* | | 63 | 62 | | | | | 19 | 18.06557 | | 62 |
| Reptilia | *Vipera latastei* | | 200.4 | 62 | | | | | 14 | | | 62 |
| Reptilia | *Woodworthia brunneus* | | 10.3 | 62 | | | | | 42 | | | 62 |
| Reptilia | *Xantusia riversiana* | | 32.2 | 62 | | | | | 32.9 | | | 62 |
| Reptilia | *Xantusia vigilis* | Desert night lizard | 9 | 38 | 6.7 | 62 | | | 11 | 0.531749 | 0.000666 | 320 |
| Reptilia | *Zootoca vivipara* | | 10.6 | 62 | | | | | 5 | | | 62 |

**Table S5.** Physiological data for extant mammals and reptiles.
